# GALILEO: Embodied AI scientist for autonomous therapeutic discovery in dynamic membrane systems

**DOI:** 10.64898/2026.06.10.731360

**Authors:** Nan Jiang, Rongyuan Wei, Hao Xiao, Zhenfei Yin, Xi Wang, Taoyong Cui, Ke Shao, Jian Zhou, Jia Fan, Philip Torr, Yingcheng Wu, Qiang Gao

## Abstract

Generalizable Agentic Laboratory Intelligence for Learning, Experimentation, and Optimization (GALILEO) closes the prediction-to-intervention gap in therapeutic peptide discovery. Unlike prior AI-scientist systems, GALILEO couples an Observation-Thought-Action-Summary (OTAS) reasoning loop with robotic peptide synthesis and multimodal phenotyping. Candidate peptides are retrieved from a clinically informed peptide prior (CPP) and locally edited through auditable operations. GALILEO autonomously prioritized LRRC8C and SLC25A1 branches and used wet-lab feedback to update target beliefs, sequence policies, assay choices, and mechanism hypotheses. Peptides generated under this framework blocked LRRC8C currents, perturbed osmolyte/redox homeostasis, and promoted tumor-dependent T-cell activation, while SLC25A1 peptides disrupted citrate-export metabolism, reduced extracellular acidosis, and enhanced CD8+ T-cell function. These results establish retrieval-and-editing-based physical learning as a route for auditable, experimentally grounded interventions and transferable membrane-blocker rules.

## INTRODUCTION

Scientific discovery is increasingly constrained by a prediction-to-intervention gap: hypotheses can now be generated faster than they can be converted into executable, target-dependent interventions in living systems. Large-language-model chemists, self-driving materials laboratories, mobile robotic chemists, AI-planned flow-synthesis platforms, and tool-augmented scientific language models have shown that agentic reasoning can be coupled to experimental planning, software tools, and physical automation.^1–7^ Therapeutic biology imposes a stricter requirement. The decisive learning signal is often not a literature-derived plausibility score or an in-silico confidence metric, but a phenotypic state transition in cells, organoids, immune cocultures, patient-derived tissues, or animals.

Recent AI-scientist systems have formalized important parts of this process. Co-Scientist generates, critiques, ranks, and refines scientific hypotheses for experimental verification,^8^ whereas Robin connects literature-search agents with experimental data-analysis agents to support iterative therapeutic hypothesis generation in experimental biology.^9^ In parallel, AI Scientist and data-to-paper-style platforms have demonstrated increasingly complete automation of computational or data-derived research cycles, including ideation, coding, experiment execution, result analysis, manuscript writing, and verification workflows. These systems make the hypothesis space cheaper to search, but they also expose a remaining bottleneck for therapeutic discovery: wet-laboratory evidence often remains a downstream validation step or a human-mediated bridge rather than the primary feedback signal for molecule-level intervention evolution.^10^ There is an urgent need to address this missing physical axis by asking whether agentic autonomy can close a therapeutic-intervention loop in living biological systems.

Dynamic membrane proteins provide a stringent testbed for this axis. Ion channels and solute carriers constitute clinically important target classes,^11^ but their therapeutic design remains difficult because membrane proteins are shaped by conformational plasticity, hydrophobic interfaces, oligomeric assembly, lipid context, and transport-cycle state.^12,13^ A useful intervention in these systems is often defined by a cellular state transition, such as pore blockade, osmolyte imbalance, metabolic collapse, extracellular-acidity remodeling, or immune activation, rather than by a static complex alone.

Therapeutic peptides offer a practical first modality for embodied discovery. Peptide therapeutics can be synthesized and iterated faster than many larger biological modalities while retaining translational relevance.^14,15^ Peptides have already been used to modulate difficult target classes, including receptor and ion-channel systems.^16,17^ Classical amphiphilicity, cell-penetrating behavior, and antimicrobial-pore principles provide important foundations,^18,19^ and computational protein design has advanced rapidly through protein language models, sequence-design algorithms, diffusion-based protein design, one-shot binder optimization, and AlphaFold-class interaction modeling.^20–27^ These advances do not, however, specify how an edited peptide should selectively occlude an endogenous channel pore or transporter cavity while remaining compatible with living-cell, organoid, and immune-context assays.

Here we present Generalizable Agentic Laboratory Intelligence for Learning, Experimentation, and Optimization (GALILEO) as a physically embodied AI scientist for closed-loop therapeutic peptide discovery in dynamic membrane systems. GALILEO is not a de novo peptide generator. Its molecular starting point is a clinically informed peptide prior, and its agentic action is retrieval, local editing, synthesis, quality control, phenotyping, and hypothesis updating under physical feedback. Applying GALILEO to ion-channel and solute-carrier targets, we show that living-system feedback can yield a transferable Amphiphilic Balance Grammar for membrane-protein blockade and two tumor-immunometabolic intervention programs: LRRC8C blockade, which couples osmolyte/redox stress to tumor-cell-intrinsic STING signaling and tumor-dependent T-cell activation; and SLC25A1 blockade, which collapses citrate-export metabolism, reduces extracellular acidification, and promotes antitumor immune permissiveness.

## RESULTS

### A clinically informed prior enables a physically embodied peptide-discovery loop

We first built GALILEO around the premise that therapeutic discovery requires both a cognitive layer that decides what should be learned and a physical layer that produces decision-grade data. The clinically informed peptide prior (CPP) was constructed from CPTAC peptide-spectrum-match (PSM) tumor proteomics across nine cancer types, with clinical metadata, sequence features, abundance profiles, cancer-type labels (Figure 1A; Figure S1A,B).^28–31^ An abundance-aware gated-fusion model with survival, recurrence, and malignancy heads extracted label-conditioned attention under favorable-outcome states, creating an auditable peptide-prior database rather than an unconstrained generator (Figure 1A). The training corpus included 1,093 patient tumor proteomes across 9 cancer types, matched adjacent-normal/control samples where available in the source cohorts, and more than 6.9 million unique PSMs; the final CPP contained 4,822 clinically anchored candidates with parent protein, cancer context, outcome label, sequence score, and physicochemical metadata (Figure 1A), and a CPP landscape scan confirmed broad distributions of peptide length, molecular weight, net charge, isoelectric point, GRAVY, aromaticity, and aliphatic index that preserve chemically diverse but synthesizable starting points for local editing (Figure S1H).

**Figure 1.**
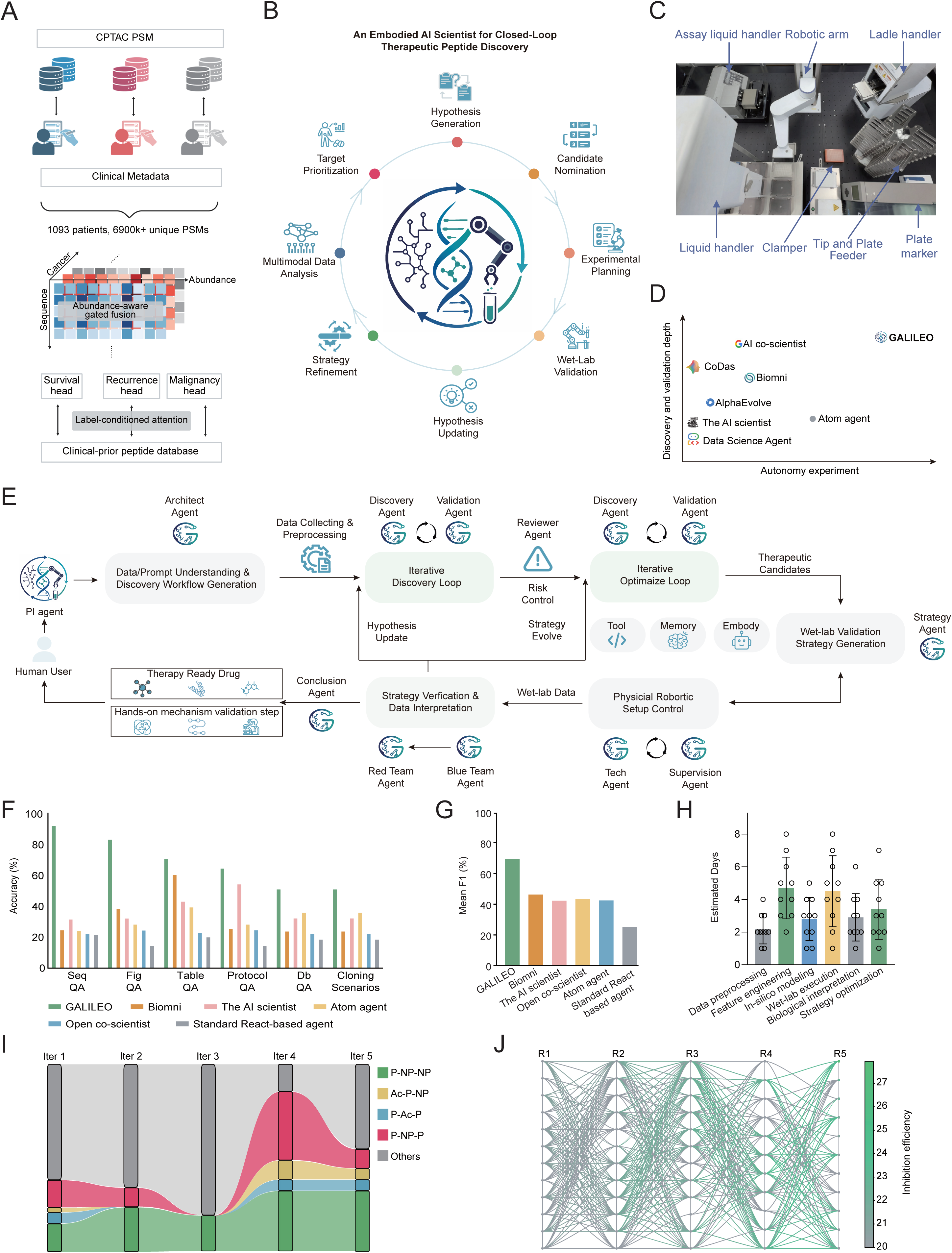
An embodied AI scientist autonomously evolves functionally active peptides. (A) Construction of the CPP. CPTAC PSM data from nine cancer types, clinical metadata, sequence features, abundance profiles, and cancer-type labels were integrated through an abundance-aware gated-fusion model with survival, recurrence, and malignancy heads. Label-conditioned attention was extracted to build the clinical-prior peptide database. The training corpus included 1,093 patient tumor proteomes, matched adjacent-normal/control samples where available in the source cohorts, and more than 6.9 million unique PSMs. (B) End-to-end GALILEO discovery workflow, linking multimodal data analysis, target prioritization, hypothesis generation, candidate nomination, experimental planning, wet-laboratory validation, hypothesis updating, and strategy refinement. Candidate peptides were retrieved from CPP and locally edited; no de novo peptide generator was used. (C) Automated wet-laboratory module landscape showing the assay liquid handler, robotic arm, labware handler, liquid handler, clamper, tip-and-plate feeder, and plate marker used for autonomous execution; the labware handler moves plates and tips, whereas the liquid handler performs reagent transfer and dosing. (D) Positioning of GALILEO relative to representative AI-scientist systems across experimental autonomy and depth of discovery/validation. Detailed scoring criteria are provided in Table S1. (E) GALILEO architecture and OTAS mapping. Observation maps to data/prompt understanding, discovery-workflow generation, and data collecting/preprocessing; Thought maps to iterative discovery and iterative optimization; Action maps to wet-lab validation and strategy generation, including automated plate-based execution when supported and hands-on validation planning when the assay exceeds the robotic module; Summary maps to strategy verification and data interpretation followed by Blue-Team/Red-Team review and hypothesis/strategy evolution. (F) Benchmark accuracy across SeqQA, FigQA, TableQA, ProtocolQA, DbQA, and cloning-scenario tasks for GALILEO and comparator agents. (G) Mean F1 across comparator agents. Mean F1 is the average harmonic mean of precision and recall across benchmark task categories. (H) Estimated human expert time required to complete the GALILEO workflow manually across data preprocessing, feature engineering, in silico modeling, wet-lab execution, biological interpretation, and strategy optimization. (I) Motif payload dynamics across five autonomous iterations, showing contraction of weak branches in iterations 2-3 followed by recovery and enrichment of productive motif families in iterations 4-5. (J) Round-resolved motif-sharing map across experimentally tested peptides. Columns denote optimization rounds, connecting lines denote shared active motifs, and line color denotes inhibition efficiency. See also Figure S1 and Tables S1 and S2.

The workflow connects multimodal data analysis, target prioritization, hypothesis generation, candidate nomination, experimental planning, wet-laboratory validation, hypothesis updating, and strategy refinement (Figure 1B; Video S1). The automated wet-laboratory module includes an assay liquid handler for reagent dispensing and plate-based assay setup, a general liquid-handling unit for dilution and transfer steps, a robotic arm, ladle handler, clamper, tip-and-plate feeder, and plate marker used for physical execution (Figure 1C). To position GALILEO within the AI-scientist landscape, we mapped representative systems by experimental autonomy and depth of discovery/validation (Figure 1D; Table S1). Whereas comparator systems emphasize hypothesis generation, literature/database reasoning, data analysis, computational research, or chemistry/materials optimization,^8,32–36^ GALILEO couples omics-informed target nomination, CPP-guided peptide retrieval and editing, robotic wet-laboratory phenotyping, feedback-driven hypothesis updating, and transferable grammar extraction. This places GALILEO on a physical-intervention axis, where living-system responses drive therapeutic molecule evolution rather than merely validating AI-ranked hypotheses.

The cognitive architecture is organized as an Observation-Thought-Action-Summary (OTAS) loop. During Observation, human prompts, data/prompt understanding, discovery-workflow generation, plate-reader outputs, Sytox Green images, CellTiter-Glo values, brightfield and fluorescence images, peptide identity, plate position, treatment concentration, incubation time, solubility or precipitation flags, and target-evidence records are converted into a structured experimental state (Figure 1E). During Thought, iterative discovery and iterative optimization loops allow the Principal Investigator (PI), Architect, Discovery, Validation, Reviewer, and Strategy agents to maintain competing target and mechanism hypotheses and to decide whether a branch should be advanced, rescued, or rejected (Figure 1E; Figure S1E). During Action, wet-lab validation and strategy generation retrieve candidates from the CPP, apply local edits when justified by the hypothesis board, rescore candidates with protein language model (PLM) or structure tools, submit SPPS and plate-based assay commands, and control robotic liquid-handling operations within the automated module (Figure 1E). When the required validation exceeds the present hardware stack - for example patch-clamp electrophysiology, Seahorse metabolic assays, patient-derived tumor-fragment testing, or in vivo studies - the same Action step produces an executable hands-on validation plan rather than claiming autonomous robotic execution (Figure 1E; Figure S1C). During Summary, strategy verification and data interpretation are reviewed by Blue-Team rule checks and Red-Team model critiques before the hypothesis board is updated (Figure 1E; Figure S1D,E).

Each automated screening round used 96-well live-cell assays seeded with 3-5 × 10³ cells per well, 10-point peptide gradients spanning 1 nM to 10 μM, and 24, 48, and 72h readouts unless a branch-specific confirmation assay required a fixed dose or expanded range (Figure S1C). The plate-based phenotypic reward integrated total cell number, Sytox Green-positive cell fraction, CellTiter-Glo viability, brightfield morphology, fluorescence intensity, peptide solubility or precipitation, HPLC-MS purity, and protocol-compliance guardrails (Figure S1C; Table S2). For multicancer prescreening, Huh-7, SNU-182, HCCLM3, and Hep3B tumor cells were evaluated with HEK293T selectivity controls, and positive branches were prioritized only when tumor inhibition was observed without matched HEK293T toxicity (Figure S2A).

Branch-level agent decisions were documented by the workflow panels, whereas the event table, hypothesis-board trajectory, and motif-evolution panels recorded sequence-policy and mechanism updates induced by feedback (Figure 1I,J; Table S2). The feedback loop changed four components of the agent state. Plate-based readouts and agent-specified hands-on validations adjusted target-branch belief; Sytox Green positivity, CellTiter-Glo viability, brightfield morphology, solubility, and QC adjusted motif-policy weights; ambiguous phenotypes triggered orthogonal validation plans such as patch clamp, organoid negative controls, siRNA-mediated target silencing, acetate rescue, Seahorse analysis, patient-derived tumor-fragment/T-cell coculture, or in vivo testing; and the hypothesis board moved from generic cytotoxicity or membrane disruption toward transport-linked stress hypotheses that were tested in the target-specific sections below (Figure 1I,J; Table S2).

Across SeqQA, FigQA, TableQA, ProtocolQA, DbQA, and cloning-scenario benchmarks,^37^ GALILEO achieved the highest task accuracy among the comparator agents (Figure 1F). GALILEO also achieved the highest mean F1, calculated as the average harmonic mean of precision and recall across benchmark answer categories (Figure 1G).^38^ Human-expert time modeling estimated that the integrated workflow reduced the days required for data preprocessing, feature engineering, in silico modeling, wet-lab execution, biological interpretation, and strategy optimization (Figure 1H). These benchmarks do not by themselves establish biological discovery; they show that the cognitive layer can coordinate heterogeneous evidence types into executable laboratory decisions.

A cognitive ablation control distinguished GALILEO from a conventional automation stack. Removing individual OTAS components reduced SciGym and LAB-Bench performance, indicating that Observation, Thought, Action, and Summary are functional coordination modules rather than interchangeable interface labels (Figure S1D). Thus, removing the reasoning layer does not merely slow the platform; it changes whether heterogeneous evidence can be converted into a valid scientific action.

Across five autonomous peptide-optimization rounds, the hypothesis-board audit captured a staged abandon-and-recover trajectory rather than monotonic enrichment (Figure 1I; Figure S1E,F; Table S2). Iteration 1 established a heterogeneous CPP-derived motif baseline; iterations 2-3 pruned weak branches and contracted motif payloads as poorly supported organizations were rejected; iteration 4 reopened previously abandoned acidic, polar, and nonpolar organizations under stronger pore-engagement hypotheses; and iteration 5 consolidated recovered Ac-P-NP and P-NP-P architectures into shared active modules across experimentally tested peptides (Figure 1I,J; Figure S1E-G). This recovery was reflected in a denser motif-sharing network, with line intensity tracking inhibition efficiency (Figure 1J). Importantly, the trajectory occurred within a chemically broad but experimentally tractable CPP landscape, as peptide length, molecular weight, net charge, isoelectric point, grand average of hydropathy (GRAVY), aromaticity, and aliphatic index all showed broad distributions compatible with local editing and synthesis (Figure S1H). Together, these data established the operating premise for the rest of the study: GALILEO can transform sparse wet-laboratory evidence into auditable design rules rather than simply ranking a static peptide list.

### Closed-loop evolution reveals an orthogonal and transferable membrane-blocker grammar

Before introducing the target identities of the retained biological campaigns, we first summarize the blocker-design grammar distilled from completed closed-loop wet-laboratory iterations and stress-tested in a larger virtual benchmark. This section is therefore a cross-branch synthesis of physical-feedback-derived rules rather than a target-selection section or a purely a priori design framework; branch-specific target-ranking evidence, target identities, and mechanisms are introduced in the subsequent biological sections (Figure 2).

**Figure 2.**
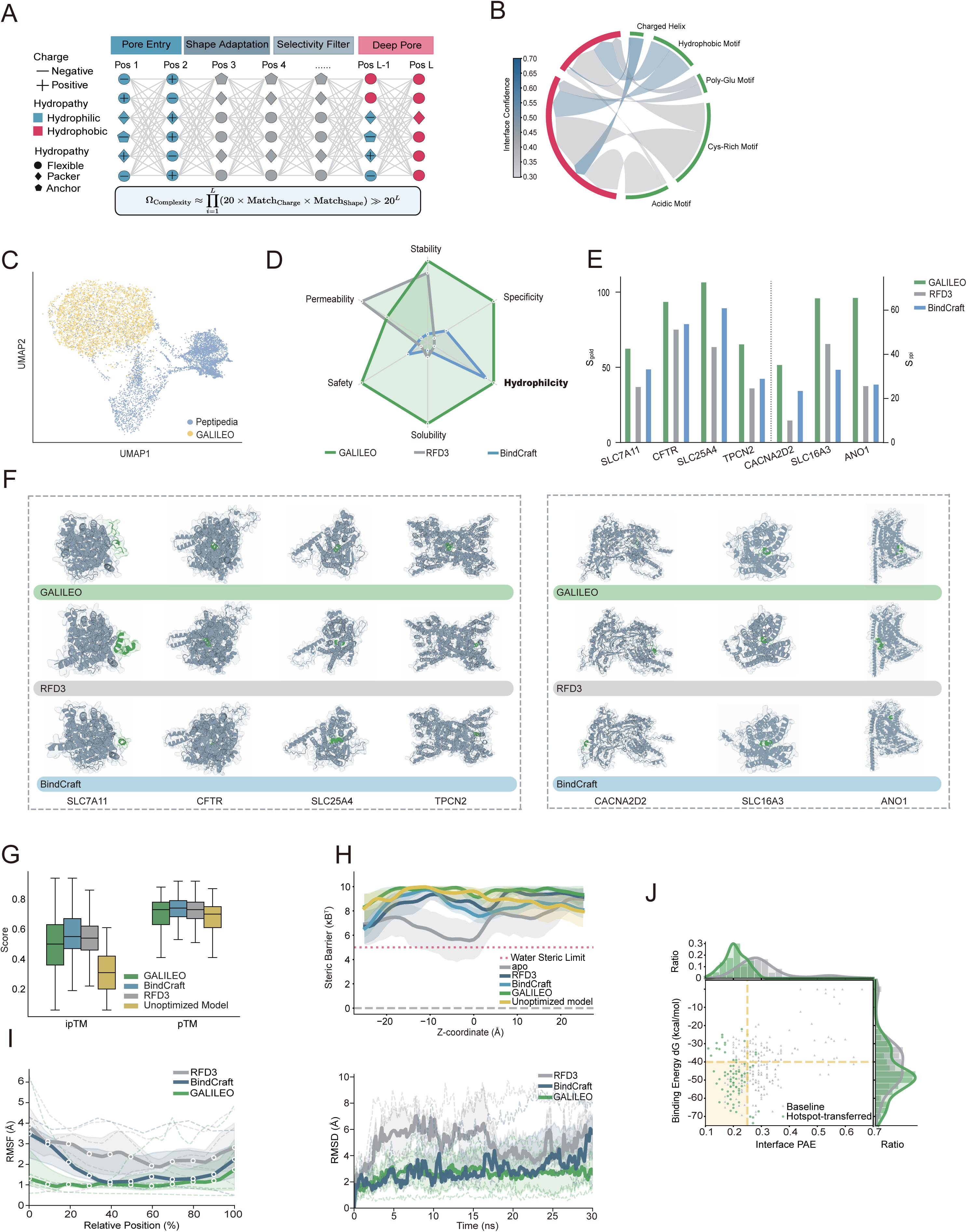
Closed-loop embodied evolution accesses an orthogonal, biophysically grounded peptide space. (A) Schematic of the target-aware blocker-design search space. For a peptide of length L, Nseq(L)=20^L. The annotated design manifold was rationally calculated as Nconstraint(L,T)=20^L × ∏i∏r |Qi,r(T)|, where Qi,r(T) represents target-conditioned categorical requirements for charge, hydropathy, flexibility, packing, and anchoring across pore entry, shape adaptation, selectivity-filter engagement, and deep-pore binding. The panel visualizes a rational design space and not an experimentally synthesized exhaustive library. (B) ABG learned during autonomous optimization. Colored sectors denote recurrent motif classes; ribbons connect motif pairings recovered in successful designs; ribbon shading denotes interface-confidence weighting. (C) UMAP projection of GALILEO-optimized peptides relative to Peptipedia-annotated reference peptides. (D) Multidimensional developability profiles of GALILEO, RFdiffusion3, and BindCraft outputs. Scores were computed with the PepVerse/PeptiVerse peptide-property scoring suite and summarized across stability, specificity, hydrophilicity, solubility, safety, and permeability axes. (E) Target-level benchmark across seven membrane-protein targets. For known-reference targets (SLC7A11, CFTR, SLC25A4, and TPCN2), blocker recovery was scored as Sgold = 100 × (0.25 Iinterface + 0.25 Rcontact + 0.20 Ooccupancy + 0.15 Ccentroid + 0.15 Mmechanism). For no-reference targets (CACNA2D2, SLC16A3, and ANO1), blocker plausibility was scored as Sppi = 100 × (0.20 BSAnorm + 0.15 Emultichain + 0.20 Aphyschem + 0.25 Ocore/pore + 0.15 Bbarrier - 0.05 Pclash). Bars show target-level means across eight AlphaFold3 rebuilds using random seeds 1-8. Paired two-sided t tests were performed on target-level AlphaFold3-averaged scores. (F) Representative rebuilt peptide-target complexes generated by GALILEO, RFdiffusion3, and BindCraft under matched computational budgets. (G) AlphaFold3-derived ipTM and pTM distributions across the membrane-target benchmark, including trained GALILEO outputs, RFdiffusion3, BindCraft, and unoptimized-model controls. (H) Steric-barrier profiles along the permeation axis for difficult targets, compared with apo, unoptimized, RFdiffusion3, BindCraft, and GALILEO models. The dashed line denotes the water steric limit. (I) Molecular-dynamics stability metrics for membrane-embedded peptide-target complexes; left, interface RMSF; right, complex RMSD across a 30-ns simulation window. (J) Interface PAE and binding-energy distributions before and after hotspot transfer of the GALILEO-derived grammar into external design workflows. See also Figure S2.

For a peptide of length L, the naive sequence space begins at 20^L possible amino-acid strings (Figure 2A). The blocker-design landscape is larger than this sequence count because each position is also assigned target-conditioned property states for charge, hydropathy, flexibility, packing, and anchoring across pore entry, shape adaptation, selectivity-filter engagement, and deep-pore binding (Figure 2A). This sequence-property-geometry manifold was rationally calculated from amino acid choices and discrete property-state assignments; it was not an experimentally synthesized exhaustive library. GALILEO differs from a self-driving laboratory that accelerates a finite combinatorial screen because physical feedback from a small tested subset can change which sequence branches are retrieved, locally edited, discarded, or reopened in the next round.

Across the retained wet-laboratory branches, closed-loop physical iteration converged on a compact blocker-design rule set that we term the Amphiphilic Balance Grammar (ABG). ABG is not generic antimicrobial or cell-penetrating amphiphilicity; rather, it is a physical-feedback-derived grammar in which acidic motifs, Cys-rich motifs, poly-Glu segments, hydrophobic motifs, and charged helices are recombined to balance pore entry with selective steric occlusion (Figure 2B). Amino-acid-pair enrichment analysis resolved this grammar into branch-conditioned, target-aware pair features that were retained after physical iteration and stress-tested through virtual iteration.

The GALILEO-edited candidates from which ABG was distilled occupied a sequence-space region orthogonal to Peptipedia-annotated reference peptides while retaining favorable developability profiles across stability, specificity, hydrophilicity, solubility, safety, and permeability (Figure 2C,D).^39^ These data position GALILEO-edited peptides in a distinct but developable peptide manifold, rather than in a novelty-only region. We benchmarked the rule-guided retrieval-and-editing strategy against RFdiffusion3 and BindCraft across seven tumor-contextualized membrane-protein targets selected to connect the CPTAC/TCGA-informed CPP search space with structurally diverse blockade tasks: SLC7A11/xCT for cystine import, redox buffering, glutathione metabolism, and ferroptosis resistance^39^; CFTR for cancer-associated epithelial chloride-conductance regulation^40^; SLC25A4/ANT1 for mitochondrial ADP/ATP carrier exchange and cancer-relevant mitochondrial metabolism^41^; TPCN2/TPC2 for endolysosomal cation signaling linked to invasive cancer-cell migration^42^; CACNA2D2 for calcium-channel auxiliary-subunit biology linked to tumor-cell proliferation and angiogenesis^43^; SLC16A3/MCT4 for lactate export, glycolytic adaptation, and tumor acidosis^44^; and ANO1/TMEM16A for calcium-activated chloride-channel signaling associated with tumor growth and invasion^45^ (Figure 2E,F). RFdiffusion3 represents a de novo backbone-diffusion, inverse-folding, and structure-filtering workflow.^25^ BindCraft represents structure-model-guided backpropagation and greedy binder optimization.^40^ All top complexes were rebuilt and rescored uniformly with AlphaFold3 across eight random seeds so that the comparison evaluated pipelines rather than isolated model seeds (Figure 2E,F).^27^

For known-reference targets (SLC7A11, CFTR, SLC25A4, and TPCN2), prior studies provided a literature-supported inhibitory region, ligand-bound pocket, or pharmacological reference, so GALILEO was evaluated by gold-reference consistency (S_gold_), which measures recovery of the relevant inhibitory interface. For no-reference targets, meaning targets without highly selective reference blockers suitable for geometry recovery (CACNA2D2, SLC16A3, and ANO1), GALILEO achieved higher intrinsic blocker plausibility, defined as S_ppi_ (Figure 2E). Under the matched evaluation pipeline, GALILEO-derived blockers more frequently occupied pore-facing or cavity-facing positions, formed broader steric barriers, and maintained target-compatible amphiphilic interfaces, indicating blocker-like geometry rather than generic surface binding (Figure 2F).

The benchmark was then expanded to 65 tumor-relevant dynamic membrane targets nominated by intersecting CPTAC/TCGA tumor-versus-normal expression, membrane localization, cancer dependency or immune-metabolic relevance, and practical peptide-blockade feasibility filters, comprising 49 SLC or SLCO transporters and 16 ion-channel or channel-associated proteins (Figure S2C). These candidates were used as a virtual-only stress test of whether the physical-branch-derived grammar generalized across tumor membrane-transport targets rather than as additional experimentally validated programs. They then underwent five rounds of internal GALILEO virtual self-iteration using the same ABG constraints and scoring parameters learned from the physical branches (Figure S2A-D). At the level of conventional AlphaFold3 confidence, GALILEO, RFdiffusion3, and BindCraft produced broadly comparable ipTM and pTM distributions after uniform rebuilding (Figure 2G; Figure S2C).

Static confidence did not fully capture blocker quality. Unoptimized models and baseline workflows could yield plausible confidence distributions, but for difficult targets they failed to generate the steric-barrier profiles and membrane-compatible geometries observed after GALILEO’s physical-learning-derived constraints were applied (Figure 2G,H; Figure S2D). Molecular-dynamics simulations in membrane environments further showed lower interfacial fluctuation and reduced complex drift for GALILEO-derived blockers, as quantified by interface RMSF and complex RMSD (Figure 2I).

To test transferability, ABG motifs discovered from the physical optimization branches were redeployed as hotspot priors in RFdiffusion3 and BindCraft. Hotspot transfer shifted baseline outputs toward lower interface predicted aligned error and more favorable binding-energy distributions relative to unconstrained runs (Figure 2J). Thus, GALILEO did not simply identify better sequences within its retained wet-lab branches; it distilled a portable intervention prior from physical feedback and used it to partially rescue blind spots in external generative design workflows (Figure 2J).

### GALILEO-nominated LRRC8C blockade perturbs osmolyte homeostasis and innate-stress signaling

To test whether GALILEO could convert multi-omics observations into an executable channel-blockade hypothesis, we prompted the agent to identify an actionable ion-channel target with broad anticancer relevance and limited pharmacological coverage. The in-house multi-cancer organoid membrane-proteomic atlas became the decision state: across BRCA, COAD, STAD, LIHC, LUNG, and PAAD organoid datasets (104 cancer organoids in total), the agent identified 35 recurrent ion-channel candidates and ranked them by expression breadth, tumor-versus-normal differential signal, druggable-pore plausibility, literature support, immune-context relevance, and unmet inhibitor need (Figure 3A; Figure S3A-C; Table S3). This decision workflow nominated LRRC8C because it combined organoid-detectable expression, LRRC8-family context, tumor-associated proliferation evidence, immune-exhaustion associations, and limited availability of selective pharmacologic blockers (Figure 3B-E; Figure S3D-F).

**Figure 3.**
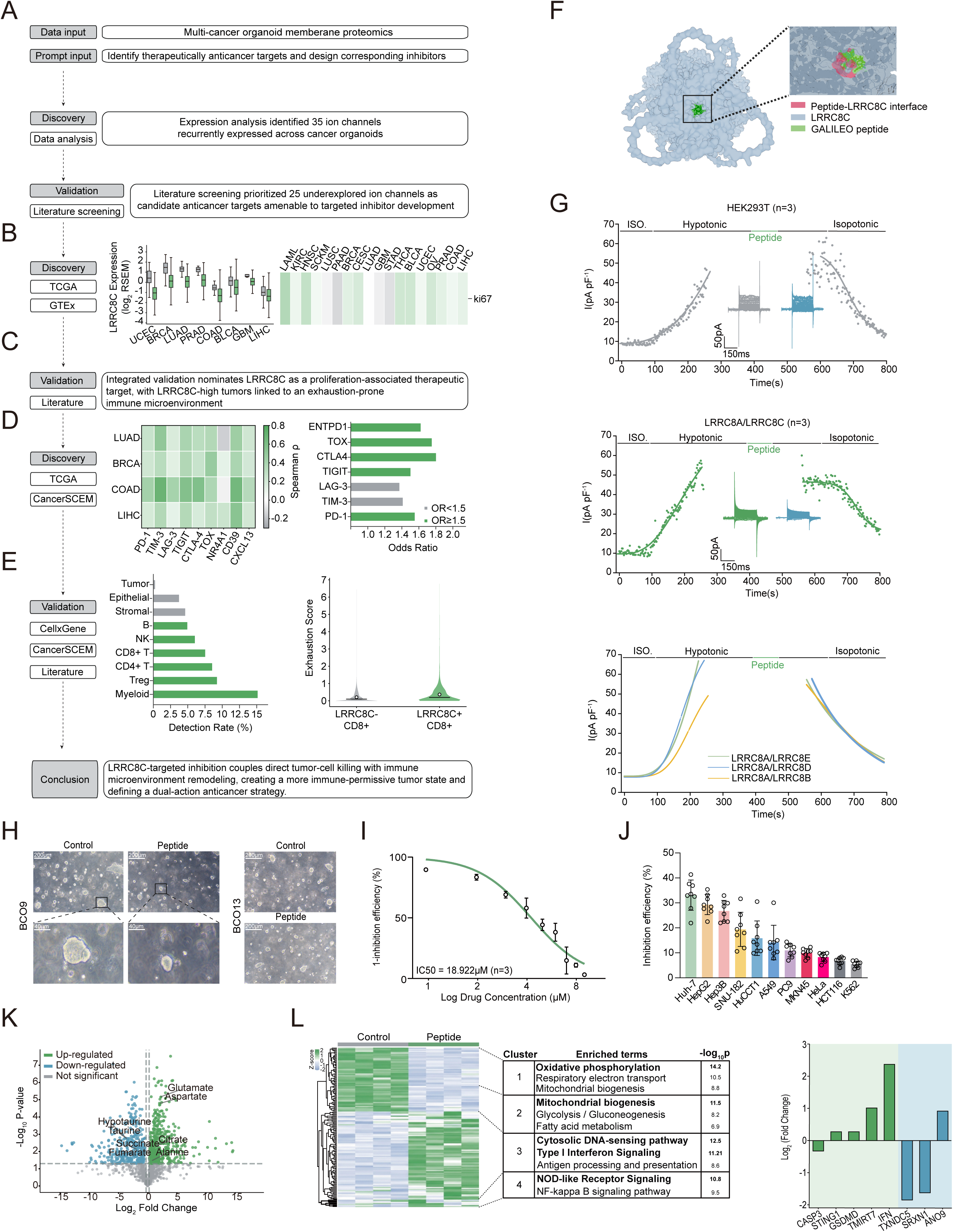
AI-designed LRRC8C blockade perturbs osmolyte homeostasis and innate-stress signaling. (A) GALILEO workflow for ion-channel target discovery from the multi-cancer organoid membrane-proteomic atlas, with literature-supported prioritization summarized in Table S3. (B) LRRC8C expression across tumor types (left, TCGA) and association with proliferative context (right). (C) Validation summary nominating LRRC8C as a therapeutically actionable ion-channel target. (D) Associations between LRRC8C and exhaustion-related features; left, correlations between LRRC8C and exhaustion markers in TCGA tumors; right, CancerSCEM-wide association between LRRC8C and T-cell exhaustion markers. (E) Single-cell analysis across 22 CancerSCEM cancer types; left, LRRC8C detection rates across tumor-microenvironment compartments; right, exhaustion scores of LRRC8C-positive and LRRC8C-negative CD8+ T cells. Exhaustion score was defined as the number of detected genes (count > 0) among PDCD1, HAVCR2, LAG3, TIGIT, CTLA4, TOX, and ENTPD1 per CD8+ T cell. (F) Structural model of GALILEO-LRC bound at a pore-facing LRRC8C interface. (G) Whole-cell patch-clamp recordings showing selective inhibition of LRRC8A/LRRC8C currents by GALILEO-LRC; top, parental HEK293T cells with minimal LRRC8C expression; middle, HEK293T cells cotransfected with LRRC8A and LRRC8C (1:1); bottom, HEK293T cells cotransfected with LRRC8A and LRRC8E, LRRC8D, or LRRC8B. (H) Representative brightfield images of LRRC8C-high and LRRC8C-low patient-derived breast cancer organoids after 72 h PBS/scrambled-peptide control or GALILEO-LRC treatment. Scale bars, 200 μm; high-magnification views, 40 μm. (I) Dose-response curve of GALILEO-LRC in LRRC8C-high patient-derived breast cancer organoids (IC50 = 18.922 μM). (J) Seventy-two-hour growth inhibition across multicancer cell lines Huh-7, HepG2, Hep3B, SNU-182, HuCCT1, A549, PC9, MKN45, HeLa, HCT116, and K562. (K) Untargeted metabolomic comparison of GALILEO-LRC-treated versus matched PBS/scrambled-peptide control Huh-7 cells, highlighting depletion of taurine, hypotaurine, and related osmolyte/redox metabolites. (L) Transcriptomic heatmap, cluster enrichment, and selected-gene changes in Huh-7 cells showing activation of cytosolic DNA-sensing, type I interferon, and NOD-like receptor programs downstream of LRRC8C blockade relative to matched PBS/scrambled-peptide controls. Data are mean ± s.d. in I and J. n = 3 independent experiments (G and I), 5 random fields per condition for representative images in H, 8 independent measurements (J), and 4 independent biological samples per group (K and L). P values were determined by unpaired two-tailed Student’s t test. ns, not significant; *P < 0.05; **P < 0.01; ***P < 0.001; ****P < 0.0001. See also Figure S3 and Table S3.

LRRC8C is a component of volume-regulated anion-channel heteromers that mediate subunit-dependent osmolyte transport and ionic-strength sensing.^41–43^ LRRC8-family channels can also participate in cyclic-dinucleotide transport, connecting channel biology to STING signaling.^49,50^ Public single-cell RNA-seq data analyses showed LRRC8C expression in both tumor and immune compartments, with pronounced signal in monocyte and T-cell populations (Figure 3D,E; Figure S3D,E). LRRC8C-positive CD8+ T cells showed higher exhaustion scores than LRRC8C-negative CD8+ T cells across cancer contexts (Figure 3E; Figure S3F). These observations aligned with prior evidence that LRRC8C can suppress T-cell function through cyclic-dinucleotide and STING-p53 signaling.^44,45^ GALILEO therefore treated LRRC8C as a tumor-immune dual-context target rather than as a tumor-cell-only marker and chose a CPP-derived peptide-blockade branch rather than a small-molecule branch because selective LRRC8C pharmacology remains limited (Figure 3D,E). Within this peptide branch, the agent retrieved a CPP-supported parent sequence and proposed local edits to create GALILEO-LRC under an emerging pore-occlusion hypothesis that was later distilled into ABG.

Structural modeling placed GALILEO-LRC at a pore-facing LRRC8C interface consistent with direct occlusion (Figure 3F). Because patch-clamp electrophysiology is outside the present robotic wet-lab module, GALILEO specified it as a hands-on validation experiment rather than an autonomous action. In HEK293T cells expressing LRRC8A/LRRC8C under hypotonic stimulation, GALILEO-LRC suppressed LRRC8A/LRRC8C currents but did not comparably inhibit parental HEK293T cells or LRRC8A/LRRC8B, LRRC8A/LRRC8D, or LRRC8A/LRRC8E containing heteromers (Figure 3G). This validation converted the structural prediction into a retained on-target mechanism.

Organoid validation shifted the branch away from nonspecific membrane toxicity and toward LRRC8C-context-dependent tumor vulnerability. GALILEO-LRC induced architectural collapse in LRRC8C-high patient-derived breast cancer organoids but caused minimal disruption in LRRC8C-low patient-derived breast cancer organoids under the same treatment protocol (Figure 3H; Figure S3G). Dose-response profiling in LRRC8C-high patient-derived breast cancer organoids yielded an IC50 of 18.922 μM in the confirmation assay (Figure 3I). A broader multicancer panel showed growth suppression across Huh-7, HepG2, Hep3B, and SNU-182 hepatocellular carcinoma cells, HuCCT1 cholangiocarcinoma cells, A549 and PC9 lung cancer cells, MKN45 gastric cancer cells, HeLa cervical cancer cells, HCT116 colorectal cancer cells, and K562 hematologic tumor cells, consistent with activity in LRRC8C-relevant cellular contexts (Figure 3J). Because LRRC8C was also detected in immune compartments, GALILEO incorporated direct immune-cell safety checks before advancing the branch: GALILEO-LRC showed negligible toxicity in Jurkat cells, and the subsequent hands-on validation explicitly tested primary T-cell activation and cell death in the absence of tumor fragments.

Because LRRC8C-containing VRAC complexes participate in osmolyte transport, GALILEO nominated osmolyte and stress-signaling assays as hands-on mechanism validation. Untargeted metabolomics in GALILEO-LRC-treated Huh-7 cells, analyzed against matched scrambled-peptide controls, revealed depletion of taurine, hypotaurine, and related osmolyte/redox metabolites (Figure 3K). Pathway enrichment highlighted taurine/hypotaurine metabolism together with TCA-cycle-associated and amino-acid-associated programs (Figure S3H). Transcriptomic profiling in GALILEO-LRC-treated Huh-7 cells, benchmarked against scrambled-peptide controls, identified activation of cytosolic DNA-sensing, type I interferon, NOD-like receptor, antigen-presentation, and NF-kappaB-associated programs (Figure 3L). The same transcriptomic analysis showed increased STING1, GSDMD, IFN, ANO9, and TMIRT7 and decreased CASP3, TXNDC5, and SRXN1 after LRRC8C blockade (Figure 3L). Together, the LRRC8C campaign connects agent-nominated channel blockade to osmolyte restriction, redox stress, and innate-stress signaling.

### GALILEO-LRC converts channel blockade into STING-linked immunogenic remodeling

Building on the GALILEO-nominated innate-stress hypothesis, the agent next specified a mechanistic validation workflow to test whether the retained LRRC8C mechanism propagated into immunogenic remodeling.^46^ In Huh-7 cells, GALILEO-LRC treatment significantly increased intracellular ROS (Figure 4A). The same treatment also caused loss of mitochondrial membrane potential measured by TMRE staining in Huh-7 cells (Figure 4B). Mitochondrial stress can engage cGAS-STING signaling,^47,48^ and LRRC8 channels have been implicated in cGAMP transport.^49–51^ Consistent with this connection, GALILEO-LRC treatment decreased extracellular cGAMP and increased intracellular cGAMP in paired Huh-7 fractions (Figure 4C). Unless otherwise stated, these mechanism assays used matched sequence-scrambled peptide controls. Together, the Huh-7 ROS, TMRE, and cGAMP measurements support a working model in which LRRC8C blockade restricts osmolyte and cyclic-dinucleotide flux, amplifies mitochondrial stress, and activates tumor-cell-intrinsic STING signaling (Figure 4D). Dose-dependent STING phosphorylation was then assayed in HepG2 cells as an orthogonal hepatocellular carcinoma context selected for robust STING immunoblot signal, whereas Huh-7 was retained for ROS, TMRE, and cGAMP continuity; this provided pathway-level confirmation of the model (Figure 4E).

**Figure 4.**
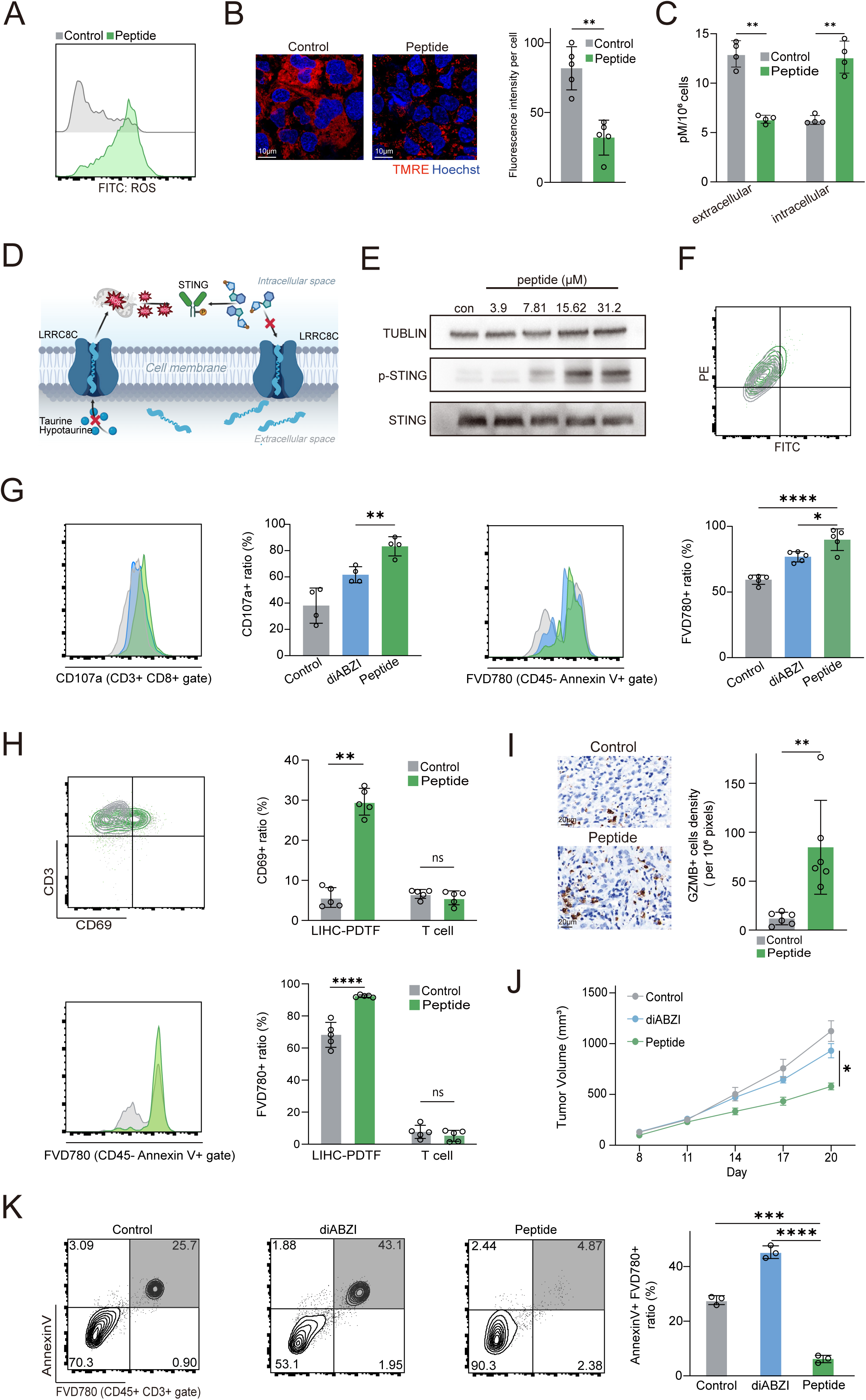
GALILEO-LRC orchestrates STING-dependent immunogenic cell death and tumor microenvironment remodeling. (A) ROS in Huh-7 cells after GALILEO-LRC treatment; representative FITC histogram. (B) TMRE/Hoechst staining in Huh-7 cells after GALILEO-LRC treatment; left, representative fields; right, quantification of TMRE fluorescence intensity per cell. Scale bars, 10 μm. (C) ELISA measurements of extracellular and intracellular cGAMP in Huh-7 cells after peptide treatment. (D) Working model linking LRRC8C blockade to osmolyte restriction, oxidative/mitochondrial stress, cGAMP redistribution, and STING activation. (E) Immunoblot of p-STING and total STING in HepG2 cells, used as an independent hepatic tumor background with robust STING immunoblot signal, after treatment with increasing concentrations of GALILEO-LRC (3.9, 7.81, 15.62, and 31.2 μM). Tubulin was used as loading control. (F) JC-1 analysis in LRRC8C-silenced Huh-7 cells; representative flow plot; righ. (G) CD107a flow cytometry and tumor-cell death readout in a 48h HLA-matched Huh-7/primary T-cell mixed-culture assay; histograms and quantification show CD107a expression within the CD3+CD8+ gate and tumor-cell death under PBS/scrambled-peptide control, diABZI, and GALILEO-LRC conditions. (H) CD3+CD69+ T-cell activation, tumor-fragment death, and isolated T-cell Annexin V/FVD780 death control in LIHC-PDTF/T-cell cocultures and isolated T-cell controls; left, representative flow plots; right, quantification. (I) GZMB-positive cell density in peptide-treated LIHC-PDTF tissues; left, representative fields; right, per-field quantification. Scale bars, 20 μm. (J) Tumor growth in an immunocompetent Hepa1-6 hepatocellular carcinoma subcutaneous tumor model treated with control, diABZI, or GALILEO-LRC; intraperitoneal treatment began on day 8. (K) Annexin V/FVD780 analysis of T-cell death under control, diABZI, and GALILEO-LRC conditions; left, representative flow plots; right, quantification of Annexin V+FVD780+ cells. Data are mean ± s.d. (A-C, F-I, and K) or mean ± s.e.m. (J). n = 4 independent experiments (A and C), 5 independent experiments or biological replicates (B, F, G, and I), 6 independent biological samples (H), 5 mice per group (J), and 3 independent experiments (K). P values for non-animal panels were determined by unpaired two-tailed Student’s t test. ns, not significant; *P < 0.05; **P < 0.01; ***P < 0.001; ****P < 0.0001. Statistical analysis of the mouse experiment in J is described in STAR Methods.

Target dependence was then tested by LRRC8C silencing. In LRRC8C-silenced Huh-7 cells, exposure to GALILEO-LRC no longer produced an additional JC-1 shift, indicating that mitochondrial depolarization required the intended channel target rather than nonspecific peptide toxicity (Figure 4F). The immune consequence of this tumor-cell-intrinsic stress program was then evaluated in a 48-h mixed culture of Huh-7 cells and HLA-matched primary human T cells. GALILEO-LRC treatment increased the CD107a-positive fraction within the CD3+CD8+ gate and, in the paired tumor-cell gate, increased the FVD780-positive tumor-cell death fraction from 59.42% in control cultures to 76.88% with diABZI (a direct STING agonist) and 89.88% with GALILEO-LRC. These paired T-cell and tumor-cell readouts indicate enrichment of a cytotoxic CD8+ T-cell phenotype with measurable tumor killing under tumor-contact conditions (Figure 4G).

Patient-derived tumor-fragment experiments provided a tissue-level test. In LIHC-PDTF cocultured with paired peripheral-blood-derived T cells, GALILEO-LRC treatment significantly expanded the CD3+CD69+ T-cell fraction and increased tumor cell death (Figure 4H). In contrast, isolated T cells maintained without tumor fragments did not show increased CD69 or Annexin V/FVD780-defined cell death after peptide treatment, supporting tumor-dependent immune activation without direct T-cell toxicity (Figure 4H). In liver hepatocellular carcinoma patient-derived tumor fragments (LIHC-PDTFs), GALILEO-LRC treatment significantly increased GZMB-positive cell density (Figure 4I).

Finally, in immunocompetent mice bearing subcutaneous Hepa1-6 tumors, GALILEO-LRC treatment suppressed tumor growth more effectively than diABZI (Figure 4J), and treatment was accompanied by stable body-weight without overt toxicity during the dosing window. Direct STING activation can support antitumor immunity, but excessive or cell-intrinsic STING signaling can also compromise T-cell viability or fitness.^52–55^ In our system, diABZI increased the density of Annexin V+FVD780+ T-cells, indicating treatment induced immune cell death, whereas GALILEO-LRC treatment preserved the effector T-cell pool while enhancing tumor-dependent immune activation (Figure 4K). Together, the ROS, mitochondrial, cGAMP, p-STING, LRRC8C-silencing, coculture, tumor-fragment, and mouse data verify the biological mechanism proposed by the GALILEO decision workflow: LRRC8C blockade is not merely cytotoxic, but a tumor-cell-intrinsic stress program that can be coupled to tumor-dependent immune activation and single-agent tumor suppression (Figure 3A-E; Figure 4A-K). Rational combinations with PD-1 blockade should be tested in future studies rather than inferred from the present dataset.

### SLC25A1-targeting peptide blockade collapses mitochondrial citrate-export metabolism

In parallel with the LRRC8C branch, GALILEO launched an SLC-transporter branch to test whether the same decision loop could generalize from ion-channel blockade to transporter blockade. The second agent objective was to identify an SLC-family target with cross-cancer therapeutic relevance and to propose an inhibitor strategy. The agent assembled a staged decision workflow composed of TCGA/GTEx discovery, GEO/CPTAC validation, immune-infiltration analysis, literature screening, and translational-gap assessment (Figure 5A). Within this workflow, SLC25A1 was prioritized over SLC7A11, SLC16A1, SLC25A51, SLC1A5, and SLC2A1 because it combined functional uniqueness, cross-cancer overexpression, prognostic relevance, druggability potential, stemness and therapy-resistance associations, and immune-modulatory relevance (Figure 5B-D).

**Figure 5.**
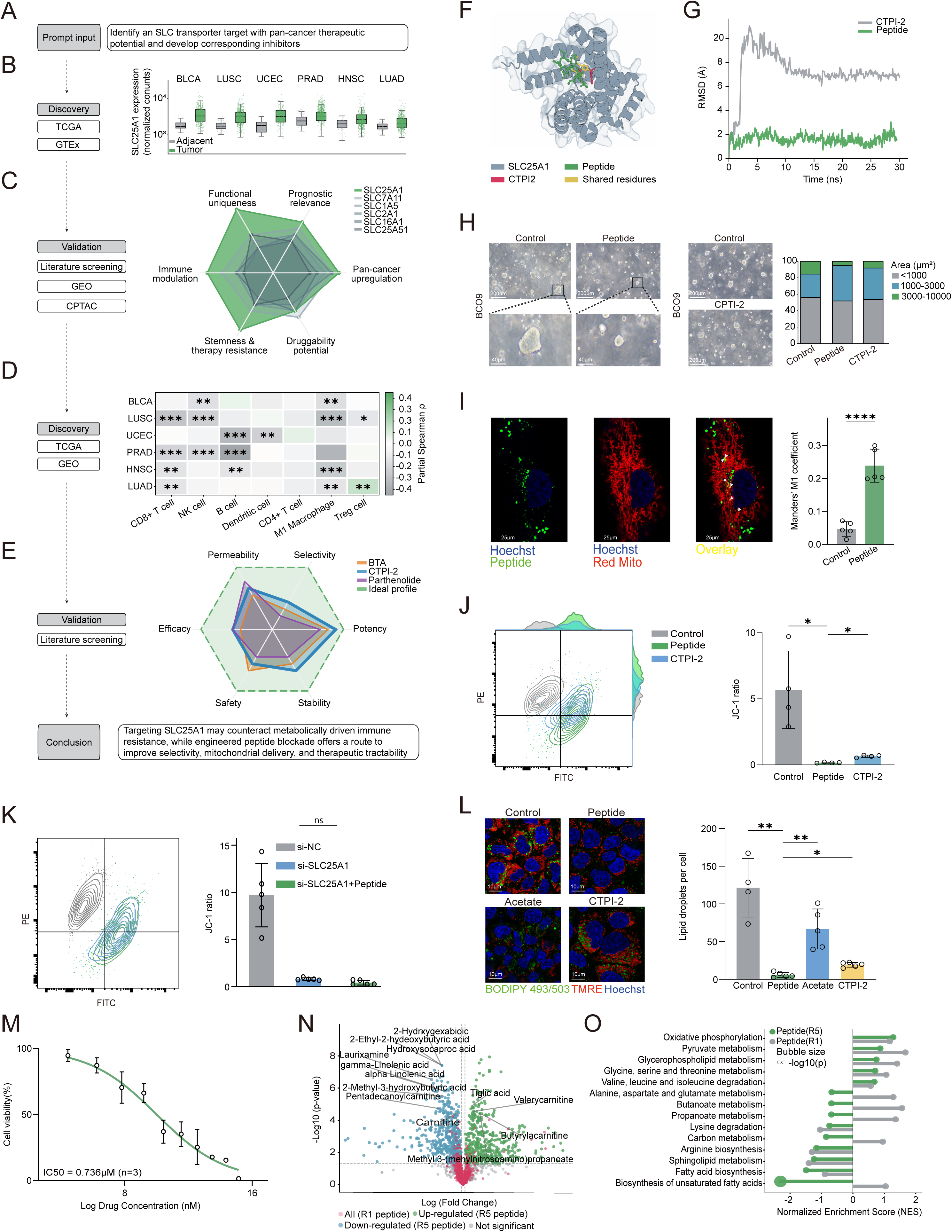
GALILEO-designed SLC25A1 blockade induces catastrophic mitochondrial collapse and metabolic reprogramming. (A) GALILEO-driven target-discovery workflow for SLC-family prioritization, integrating TCGA and GTEx discovery analyses with GEO, CPTAC, and literature validation. (B) Paired tumor-versus-adjacent expression of SLC25A1 across BLCA, LUSC, UCEC, PRAD, HNSC, and LUAD in TCGA. (C) Multi-criteria comparison of candidate SLC targets across functional uniqueness, prognostic relevance, pan-cancer overexpression, druggability potential, stemness/drug resistance, and immune modulation. (D) Spearman correlations between SLC25A1 expression and immune-cell infiltration across tumor types, computed from TCGA and GEO data using CIBERSORT. (E) Translational-gap analysis of current SLC25A1 chemotypes benchmarked against an ideal inhibitor across potency, selectivity, permeability, efficacy, toxicity, and stability. (F) Structural model positioning GALILEO-SLC and CTPI-2 within the SLC25A1 substrate-binding cavity; shared contact residues are highlighted. (G) Molecular-dynamics RMSD comparing GALILEO-SLC and CTPI-2 in a mitochondrial membrane system generated with CHARMM-GUI and simulated for 30 ns with GROMACS. (H) BCO9 breast cancer organoids after 72 h control, GALILEO-SLC, or equimolar CTPI-2 treatment; left, representative fields; right, area-distribution quantification. Scale bars, 200 μm; high-magnification views, 40 μm. (I) Mitochondrial colocalization of 5-FAM-Ahx-labeled GALILEO-SLC in cells stained with Red MitoTracker. Green, GALILEO-SLC; red, mitochondria; yellow, colocalization at the mitochondrial membrane. Scale bars, 25 μm. (J) JC-1 analysis comparing GALILEO-SLC with CTPI-2; left, representative flow plots; right, quantification of the JC-1 aggregate/monomer ratio (PE/FITC). (K) JC-1 analysis in control and siRNA-mediated SLC25A1-silenced cells; left, representative flow plots; right, quantification. (L) BODIPY/TMRE/Hoechst imaging and lipid-droplet quantification in Huh-7 cells after 72 h of single-dose treatment with GALILEO-SLC or CTPI-2, with acetate rescue; left, representative fields; right, ImageJ quantification. Scale bars, 10 μm. (M) Dose-response curve of GALILEO-SLC in SLC25A1-high BCO9 breast cancer organoids (IC50 = 736.22 nM, equivalent to 0.736 μM); equimolar CTPI-2 produced weaker organoid phenotypes in H. (N) Untargeted metabolomic profiling in Huh-7 cells comparing iteration 5 and iteration 1 peptides at matched inhibitory effect, with each peptide condition analyzed relative to its matched untreated control. (O) Pathway enrichment of the differential metabolite sets in N. Data are mean ± s.d. n = 5 random fields per condition (H), 4 independent experiments (J and L), 5 independent experiments (K), 3 independent experiments (M), and 4 independent biological samples per group (N and O). P values were determined by unpaired two-tailed Student’s t test. ns, not significant; *P < 0.05; **P < 0.01; ***P < 0.001; ****P < 0.0001. See also Figure S4.

SLC25A1 is the only known human mitochondrial citrate transporter and links mitochondrial citrate export to lipogenesis, acetyl-CoA metabolism, redox balance, and therapeutic resistance.^56–59^ Lipogenic and citrate-export dependencies provide metabolic opportunities but also translational challenges for cancer therapy.^58^ From this evidence, GALILEO identified a translational gap: available SLC25A1 chemotypes, including CTPI-2 and related small molecules, remain constrained by selectivity, mitochondrial delivery, potency, toxicity, and stability (Figure 5E; Figure S4A,B). The agent therefore chose a CPP-derived peptide-blockade branch rather than a small-molecule optimization branch (Figure 5A,E).

Structural modeling placed GALILEO-SLC within the SLC25A1 substrate-binding cavity in a pose that partially overlapped with CTPI-2, the benchmark small-molecule inhibitor, while extending beyond the small-molecule contact region (Figure 5F). Thus, GALILEO-SLC preserved shared contact residues with CTPI-2 yet created a broader interface, consistent with a distinct blocking geometry (Figure 5F). Molecular dynamics in a mitochondrial-membrane system generated with CHARMM-GUI and simulated with GROMACS showed more stable engagement and lower interfacial fluctuation for GALILEO-SLC than for CTPI-2 (Figure 5G; Figure S4C).^60^ All direct cell and organoid comparisons with CTPI-2 in this section were performed under matched molar dosing unless otherwise stated, to ensure that differences in phenotype are interpretable at the same molecular exposure.

Functional assays confirmed that the predicted interaction translated into cellular activity. In SLC25A1-high patient-derived breast cancer organoids, GALILEO-SLC or equimolar CTPI-2 was added for 72 h before organoid-size quantification from randomly sampled fields. GALILEO-SLC induced marked architectural collapse, whereas CTPI-2 produced only modest shrinkage (Figure 5H). A 5-FAM-Ahx-labeled GALILEO-SLC peptide entered cells and colocalized with MitoTracker-labeled mitochondria, supporting delivery to the organellar compartment where SLC25A1 resides (Figure 5I). GALILEO-SLC caused near-complete loss of JC-1 aggregate formation, exceeding the effect of equimolar CTPI-2, and this mitochondrial depolarization was largely abolished by siRNA-mediated SLC25A1 silencing (Figure 5J,K). Dose-response profiling in SLC25A1-high cancer organoids yielded an IC50 of 0.736 μM, together with the matched-molar CTPI-2 comparison and the reported micromolar SLC25A1-binding potency of CTPI-2 (IC50 ≈ 3.5 μM),^61^ supporting stronger phenotypic potency of the peptide blockade strategy at equivalent molecular exposure (Figure 5M).

Mechanistically, GALILEO-SLC treatment produced a collapse of mitochondrial metabolic homeostasis. In Huh-7 cells, peptide treatment reduced TMRE fluorescence and nearly eliminated cytoplasmic lipid droplets, whereas equimolar CTPI-2 produced a significantly weaker phenotype (Figure 5L; Figure S4D). Exogenous acetate partially restored lipid storage and mitochondrial polarization, supporting blockade of citrate-export-dependent lipogenic flux rather than nonspecific intracellular damage (Figure 5L; Figure S4D). Orthogonal ROS measurements in Huh-7 cells showed stronger oxidative stress with GALILEO-SLC treatment than with equimolar CTPI-2 (Figure S4E).

Mitochondrial colocalization, siRNA-mediated SLC25A1 silencing, acetate rescue, and iteration-resolved metabolomics progressively shifted the GALILEO hypothesis board from simple cavity occupancy to a GALILEO-proposed citrate-export-dependent metabolic-collapse mechanism. The agent then placed acetate rescue, mitochondrial-potential measurement, ROS quantification, and metabolomic profiling into the hands-on validation sequence. Untargeted metabolomics in Huh-7 cells comparing iteration 1 and iteration 5 peptides at matched inhibitory effect showed that the optimized peptide drove deeper perturbation of carnitine-linked and fatty-acid-related intermediates, including carnitine, valeryl carnitine, and butyryl carnitine (Figure 5N). Pathway-level analysis identified stronger suppression of oxidative phosphorylation, pyruvate metabolism, glycerophospholipid metabolism, fatty-acid biosynthesis, and related mitochondrial programs after iteration 5 peptide treatment (Figure 5O).

### SLC25A1 blockade alleviates tumor-associated acidity and enhances antitumor immune permissiveness

Because GALILEO’s hypothesis board inferred that SLC25A1-dependent citrate-export blockade could reduce tumor cell-driven acid production, the agent next asked whether this metabolic state transition could reshape tumor-immune conditions. The agent proposed that citrate-export blockade could reduce extracellular acidification, a known barrier to T-cell activity, and placed extracellular pH profiling, Seahorse extracellular acidification rate (ECAR) analysis, LIHC-PDTF test, and T-cell coculture assays into the hands-on validation queue.

Transcriptomic profiling in Huh-7 cells compared early and late GALILEO peptide iterations at matched inhibitory effect, with each treatment analyzed against its matched untreated control (Figure 6A). Iteration 5 peptide induced stronger positive enrichment of macroautophagy, autophagy, and necroptosis programs and stronger negative enrichment of glycolipid metabolism and cholesterol biosynthesis than iteration 1 (Figure 6A). Autophagic-flux imaging showed a marked increase in autophagosomes and autolysosomes after optimized peptide treatment in Huh-7 cells (Figure 6B). Immunoblotting showed dose-dependent AMPKα phosphorylation across 58.6-3,750 nM GALILEO-SLC while total AMPK remained stable, consistent with AMPK acting as an energy-stress sensor (Figure 6C).^62,63^

**Figure 6.**
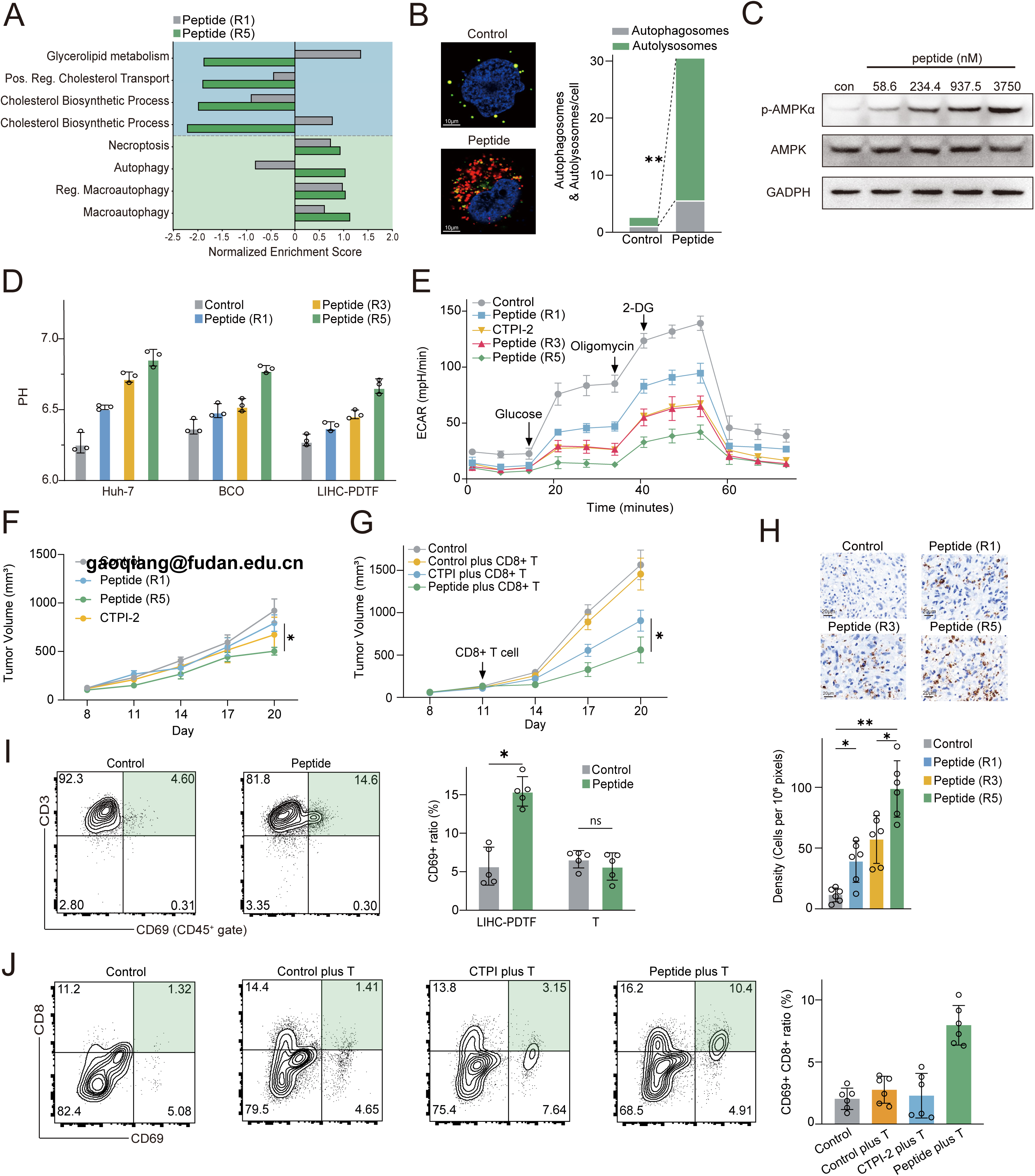
Metabolic reprogramming alleviates tumor-associated acidity and enhances antitumor immunity. (A) Transcriptomic pathway enrichment in Huh-7 cells comparing iteration 1 and iteration 5 SLC25A1-targeting peptides at matched inhibitory effect, with each peptide condition analyzed relative to its matched untreated control. (B) Autophagic-flux imaging in Huh-7 cells after peptide treatment; left, representative fields; right, quantification of autophagosomes and autolysosomes per cell. Scale bars, 10 μm. (C) Immunoblot of p-AMPKα and total AMPK in Huh-7 cells 48 h after treatment with increasing concentrations of GALILEO-SLC (58.6, 234.4, 937.5, and 3,750 nM). (D) Extracellular pH measurements in Huh-7 cells, breast cancer organoids, and LIHC-PDTF cultures across control and peptide-iteration conditions. (E) ECAR traces from a hands-on Seahorse glycolytic stress test in Huh-7 cells across treatment conditions. Glucose, oligomycin, and 2-deoxyglucose (2-DG) injections are indicated. (F) Tumor growth in Hepa1-6 hepatocellular carcinoma-bearing immunocompetent mice after control, iteration 1 peptide, iteration 5 peptide, or CTPI-2 treatment. (G) Tumor growth in vivo in a B16-OVA adoptive-transfer model using OT-1 CD8+ T cells without additional OVA antigen engineering during treatment; the arrow indicates the day of OT-1 CD8+ T-cell transfer. Groups are denoted in the manuscript as control, control plus OT-1 CD8+ T cells, CTPI-2 plus OT-1 CD8+ T cells, and GALILEO-SLC plus OT-1 CD8+ T cells to avoid ambiguity with marker-positive notation. (H) Ex vivo CD69+CD8+ T-cell flow cytometry from the adoptive-transfer experiment; left, representative flow plots; right, quantification. (I) CD3+CD69+ T-cell activation in LIHC-PDTF/T-cell cocultures and isolated T-cell controls; left, representative flow plots from the LIHC-PDTF coculture condition; right, quantification. (J) GZMB immunohistochemistry in LIHC-PDTF tissues; left, representative tumor fields; right, quantification of GZMB-positive cell density. Scale bars, 20 μm. Data are mean ± s.d. (B, D, E, H, I, and J) or mean ± s.e.m. (F and G). For A, control and peptide groups each comprised 4 independent biological samples. Quantification in B was based on 5 random fields. n = 6 independent biological samples (H and J), 5 independent experiments or biological replicates (I), 5 mice per group (F and G), 3 independent biological replicates (D), and 4 independent experiments (E). P values for non-animal panels were determined by unpaired two-tailed Student’s t test. ns, not significant; *P < 0.05; **P < 0.01; ***P < 0.001; ****P < 0.0001. Statistical analyses of the mouse experiments in F, G, H, and J are described in STAR Methods.

Extracellular acidosis is a major barrier to antitumor immunity: solid tumors commonly exhibit an extracellular pH of about 6.5-6.9, a range that suppresses T-cell activation and can blunt immunotherapy responses.^64–67^ Restoring extracellular pH toward physiological levels is therefore therapeutically attractive, although durable and tumor-selective correction remains nontrivial. We therefore asked whether SLC25A1 blockade by GALILEO-SLC could alleviate tumor cell-driven extracellular acidification.

Across Huh-7 cells, breast cancer organoids, and LIHC-PDTF cultures, successive peptide iterations progressively increased extracellular pH, with iteration 5 shifting the medium from approximately 6.2 to 6.8 in Huh-7 cells, 6.3 to 6.75 in BCO organoids, and 6.25 to 6.65 in LIHC-PDTF cultures (Figure 6D). Seahorse glycolytic stress testing in Huh-7 cells showed a stepwise decrease in ECAR, and iteration 5 produced the lowest ECAR profile; where CTPI-2 was included, comparisons were performed under matched molar exposure (Figure 6E). These pH and ECAR results are consistent with evidence that tumor-derived lactate and acidic niches suppress antitumor T-cell activity.

The in vivo relevance of this metabolic remodeling was tested in two efficacy settings. In immunocompetent mice bearing Hepa1-6 tumors, administration of iteration 5 peptide reduced tumor volume by day 20 more effectively than iteration 1 peptide or CTPI-2 (Figure 6F). In a B16-OVA adoptive-transfer model using OT-1 CD8+ T cells, without additional OVA antigen engineering during treatment, mice receiving GALILEO-SLC plus OT-1 CD8+ T cells showed stronger tumor control than mice receiving control plus OT-1 CD8+ T cells or CTPI-2 plus OT-1 CD8+ T cells (Figure 6G). Flow cytometry from the adoptive-transfer mouse experiment showed the highest CD69+CD8+ fraction in the GALILEO-SLC plus CD8+ T-cell group (Figure 6H). In LIHC-PDTF/T-cell cocultures, GALILEO-SLC increased the CD3+CD69+ T-cell fraction from 4.60% to 14.6%, whereas isolated T cells cultured without tumor fragments showed no significant activation (Figure 6I). In LIHC-PDTF tissues, immunohistochemistry showed iteration-dependent increases in GZMB-positive cell density, with the strongest staining detected after iteration 5 peptide treatment (Figure 6J). Together, these data indicate that GALILEO optimized an SLC25A1-targeting peptide that disrupts mitochondrial citrate-export metabolism, alleviates tumor-associated extracellular acidification, improves tumor control, and creates a more permissive environment for antitumor T-cell activation.

## DISCUSSION

The rapid progress of AI-for-Science has been driven by systems that retrieve evidence, generate hypotheses, rank candidates, and automate scientific reasoning. Yet therapeutic discovery faces a deeper bottleneck: biological insight becomes actionable only when converted into molecules that perturb living systems and improve through measured consequences. Our study addresses this bottleneck by defining a physical learning layer between agentic reasoning and experimental biology. GALILEO connects clinically anchored priors, local molecular editing, robotic execution, multimodal phenotyping, and recursive hypothesis updating into a closed intervention loop. Here, the wet laboratory is not a terminal validation step for computational predictions, but a source of decision-grade supervision. The LRRC8C and SLC25A1 programs therefore do not define the limits of GALILEO; they establish proof of principle that living-system response can guide therapeutic molecule evolution.

This work provides a roadmap for moving autonomous discoveries from candidate nomination to experimentally learned design logic. The Amphiphilic Balance Grammar captures this transition by linking motif organization to pore or cavity engagement, stress propagation, and tumor-immunometabolic state transitions. Many difficult targets are biologically compelling but hard to drug because their actionable states are dynamic, membrane-embedded, conformationally plastic, or poorly captured by static structural confidence.^68^ GALILEO turns these constraints into learnable features through recursive perturbation and measurement. Although developed here for ion channels and solute carriers, this principle could extend to weakly ligandable, allosteric, intracellular, or state-dependent targets whenever compact biological rewards can be measured. In such settings, AI does more than accelerate search; it converts perturbations into reusable design priors.

This concept is also relevant when model performance is limited by sparse functional labels. Tumor neoantigen prediction, for example, lacks dense paired data connecting antigen sequence, presentation, T-cell recognition, and immune function. A GALILEO-like loop could generate antigen libraries, stimulate engineered or primary T cells, measure activation, cytotoxicity, exhaustion, cytokine output, and clonotype expansion, and recycle these outcomes into antigen-prioritization models. Similar lab-in-the-loop strategies could support immune-agonist optimization, cytokine engineering, CAR/TCR tuning, synthetic-lethality screening, and patient-derived organoid testing.^69–71^ Across these applications, the decisive variable is not binding alone, but whether an intervention produces a measurable and therapeutically useful biological state transition.

The two therapeutic programs provide biological evidence for this principle without reducing GALILEO to two target discoveries. LRRC8C and SLC25A1 represent distinct membrane architectures and stress routes, yet both converged on a shared therapeutic logic: membrane-transport blockade can redirect tumor-intrinsic osmotic or metabolic vulnerability toward immune-permissive remodeling. This interpretation is bounded by prior evidence linking LRRC8-containing channels to cyclic dinucleotide transport and STING-dependent immune regulation, by reports that direct STING activation can impose context-dependent limits on T-cell fitness, and by studies identifying SLC25A1 as a mitochondrial citrate carrier supporting cancer-cell metabolic adaptation. The significance is not that GALILEO reproduced known biology, but that it linked target selection, molecular refinement, and living-system phenotypes into a recursive route for discovering intervention rules.

The current study also clarifies the boundary of physical AI. GALILEO is most scalable when experimental modules are repeatable, quantitative, and automatable, including synthesis, QC, liquid handling, imaging, and plate-based phenotyping. Specialized physiological assays, patient-derived tissue handling, and organism-level validation remain partially hands-on and define the next hardware-integration frontier. Translation will require broader pharmacokinetic, biodistribution, formulation, immunogenicity, and long-term toxicity studies, particularly for peptide modalities.^72^ The generality of the Amphiphilic Balance Grammar beyond membrane-protein blockade, peptide inhibitors, and the tumor models studied here also remains to be mapped. In summary, GALILEO establishes a mode of AI-driven therapeutic discovery in which hypotheses become molecules, molecules become perturbations, and perturbations become transferable rules learned from living-system response.

### Limitations of the Study

Several limitations define the next engineering and translational frontier for GALILEO. The current robotic layer automates synthesis, QC, liquid handling, imaging, and plate-based phenotyping, but electrophysiology, Seahorse profiling, patient-derived tissue experiments, and in vivo validation remain agent-specified hands-on modules. Scaling the platform will require robotic integration of these assays, external blinded benchmarks, and release of decision logs, protocol traces, assay metadata, and failure modes for auditability. The therapeutic programs remain preclinical; peptide translation will require systematic evaluation of protease resistance, serum stability, biodistribution, clearance, tissue and organellar delivery, immunogenicity, off-target pharmacology, formulation, manufacturability, formal toxicology, and rational combinations such as PD-1 blockade. Finally, the Amphiphilic Balance Grammar remains incompletely resolved at atomic resolution in native lipid bilayers, requiring cryo-EM, membrane reconstitution, loop-integrated structural readouts, and larger blinded target panels to define its generality.

## Abbreviations

ABG: Amphiphilic Balance Grammar
BCO: breast cancer organoid
CPP: clinically informed peptide prior
CPTAC: Clinical Proteomic Tumor Analysis Consortium
CTPI-2: citrate transport protein inhibitor 2
ECAR: extracellular acidification rate
GEO: Gene Expression Omnibus
GTEx: Genotype-Tissue Expression
GZMB: granzyme B
HPLC-MS: high-performance liquid chromatography-mass spectrometry
PDTF: patient-derived tumor fragment
OTAS: Observation-Thought-Action-Summary
PLM: protein language model
PSM: peptide-spectrum match
SPPS: solid-phase peptide synthesis
STING: stimulator of interferon genes
TCGA: The Cancer Genome Atlas
TMRE: tetramethylrhodamine ethyl ester
VRAC: volume-regulated anion channel

## ACKNOWLEDGEMENTS

This study was supported by the National Natural Science Foundation of China (82130077, 82341008, 92459301, 82441049, 92574302), the Shanghai Science and Technology Commission (No. 23JS1410200), the Fundamental and Interdisciplinary Disciplines Breakthrough Plan of the Ministry of Education of China (No. JYB2025XDXM508), the Lingang Laboratory (LGL-8888-07), the Shanghai Academy of Natural Sciences (SANS) Exploration Scholars Project, and the New Cornerstone Science Foundation.

## AUTHOR CONTRIBUTIONS

Q.G., Y.C.W., P.T., X.W., T.Y.C. and N.J. conceived the study, defined the scientific objectives and supervised the project. N.J., H.X. and Z.F.Y. designed, implemented and validated the GALILEO embodied AI scientist and the closed-loop OTAS optimization platform. Under researcher-defined objectives, constraints and validation criteria, GALILEO autonomously generated and iteratively refined the target hypotheses, peptide candidates, structural models, benchmarking decisions, experimental plans, robotic-execution commands, multimodal data analyses and model-updating decisions reported in this study. N.J. and R.Y.W. curated the clinical-proteomic, structural and public-database priors, configured the target-discovery and benchmarking tasks, and audited the computational outputs. N.J., R.Y.W. and S.K. established the automated wet-laboratory and validation workflows, monitored GALILEO-directed peptide screening and multimodal phenotyping, and performed orthogonal cell-line, organoid, PDTF/T-cell co-culture, omics, imaging, flow-cytometry, electrophysiology and mouse validation assays. N.J. and H.X. curated the source data and code, maintained the analysis pipelines, and verified statistical analysis and visualization. N.J. drafted the manuscript with input from all authors. Q.G., Y.C.W. and P.T. revised the manuscript and finalized the study. All authors discussed the results, approved the final manuscript and take responsibility for the integrity of the work. GALILEO is not listed as an author.

## DECLARATION OF INTEREST

The authors declare no competing interests.

## SUPPLEMENTAL INFORMATION

Supplemental information can be found in the submitted files.

### STAR⍰METHODS

#### KEY RESOURCES TABLE

The key resources table is available in the submitted files.

#### RESOURCE AVAILABILITY

##### Lead contact

Further information and requests for resources and reagents should be directed to and will be fulfilled by the Lead Contact, Qiang Gao.

##### Materials availability

Peptide sequences, synthesis conditions, HPLC-MS quality-control records, and cell-line or organoid metadata will be provided in Supplementary Data files or made available from the lead contact under appropriate material-transfer agreements.

##### Data and code availability

CPP construction scripts, prompt schemas, decision logs, scoring scripts, and analysis notebooks should be deposited with private review links during peer review and public or controlled-access links upon publication.

#### EXPERIMENTAL MODEL DETAILS

##### Human samples and ethics

Patient-derived breast cancer organoids, LIHC-PDTFs, paired or HLA-compatible peripheral-blood-derived T cells, and in-house hepatobiliary tumor proteomic materials were obtained from de-identified human samples under the Institutional Review Board of Zhongshan Hospital, Fudan University (No. B2023-350). Human analyses were conducted in accordance with the Declaration of Helsinki and institutional data-governance rules. Patient-derived organoid and PDTF samples were generated from surgical resection specimens with written informed consent, de-identified before downstream experiments, and analyzed only within the approved governance framework. De-identified donor blood samples used for T-cell isolation were handled under the same approved framework.

##### Cell lines

Huh-7, HepG2, Hep3B, SNU-182, HCCLM3, HuCCT1, A549, PC9, MKN45, HeLa, HCT116, K562, HEK293T, Hepa1-6, Jurkat, and B16-OVA cells were cultured at 37°C in a humidified CO2 incubator (HERAcell 150i, Thermo Fisher Scientific) with 5% CO2. Unless supplier-specific medium was required, adherent epithelial tumor cell lines were maintained in DMEM (Gibco) or RPMI-1640 (Gibco) supplemented with 10% fetal bovine serum (FBS; Gibco), 100 U ml-1 penicillin (Gibco), and 100 μg ml-1 streptomycin (Gibco); HCT116 cells were maintained in McCoy’s 5A medium (Gibco) with 10% FBS; K562 and Jurkat cells were maintained in RPMI-1640 medium (Gibco) with 10% FBS; and HEK293T cells were maintained in DMEM (Gibco) with 10% FBS. Cells were used within a low-to-moderate passage range after thawing, were routinely tested to be mycoplasma-negative, and were authenticated by STR profiling or by source documentation.

##### Patient-derived organoids

Patient-derived cancer organoids were established from surgical resection specimens from six cancer types, including breast invasive carcinoma (BRCA), colon adenocarcinoma (COAD), stomach adenocarcinoma (STAD), liver hepatocellular carcinoma (LIHC), lung adenocarcinoma (LUAD), and pancreatic adenocarcinoma (PAAD), following established patient-derived organoid principles.^73^ Organoids were cultured in growth-factor-reduced Matrigel (Corning) using cancer-type-specific organoid media and passaged every 7-10 days. BCO9 and BCO13 patient-derived breast cancer organoids were used as LRRC8C-high and LRRC8C-low contexts, respectively; BCO9 was also used as an SLC25A1-high breast cancer organoid context. LRRC8C and SLC25A1 expression status was verified by qPCR using a real-time PCR system (QuantStudio 5, Applied Biosystems/Thermo Fisher Scientific) and, where available, by protein-level or transcript-level profiling before treatment.

##### LIHC-PDTFs and PDTF/T-cell cultures

Fresh LIHC tumor tissues were processed into approximately 1-2 mm3 fragments within 2-4 h of surgical resection whenever possible, consistent with ex vivo patient-derived tumor-fragment culture workflows.^74^ Fragments were washed in cold PBS containing antibiotics, randomized into treatment groups, and cultured in RPMI-1640 medium (Gibco) or Advanced DMEM/F12-based medium (Gibco) supplemented with 10% FBS or B27/N2 supplement (Thermo Fisher Scientific) as appropriate, 1% penicillin-streptomycin (Gibco), and 10 mM HEPES (Sigma-Aldrich). For PDTF/T-cell cocultures, fragments were cultured with paired or HLA-compatible peripheral-blood-derived T cells for 72 h before flow-cytometry or immunohistochemistry readouts. Figure legends indicate n=6 LIHC-PDTF biological samples for GZMB density analysis and n=5 independent PDTF/T-cell experiments where applicable.

##### Primary human T cells and matching

Primary human T cells were isolated from fresh peripheral blood using the EasySep negative-selection workflow (STEMCELL Technologies) according to the manufacturer’s protocol. Cells were washed in PBS containing 2% FBS, counted by trypan blue exclusion using a cell counter or hemocytometer (Countess 3 FL Automated Cell Counter, Thermo Fisher Scientific; Neubauer hemocytometer, Marienfeld), and used without exogenous activation unless otherwise specified. HLA compatibility or pairing was assessed using available clinical pairing information and flow-cytometry-based matching where applicable; HLA-ABC/HLA-DR expression and CD3/CD8 purity were confirmed by flow cytometer (CytoFLEX S, Beckman Coulter) before coculture. Unless the figure legend specified otherwise, T cells were mixed with tumor cells or fragments at a conventional effector:target ratio of 5:1 to 10:1 and cultured for 72 h.

##### Animal models

Animal experiments were performed under institutional animal-care procedures, with humane endpoints applied according to approved laboratory practice. Female C57BL/6 mice, 7 weeks of age, were housed under specific pathogen-free conditions with controlled temperature, humidity, and a 12-h light/dark cycle, with free access to food and water. For the LRRC8C in vivo study, mice were subcutaneously inoculated with 2 x 10^6 Hepa1-6 cells. When tumors reached 50-100 mm3, mice were randomized and treated with vehicle, GALILEO peptide (40 μg per dose), or diABZI (40 μg per dose), a systemic aminobenzimidazole STING agonist used as an active immune comparator, administered every 3 days for four doses; tumor volume was measured every 2 days. For the SLC25A1 iteration model, C57BL/6 mice bearing Hepa1-6 tumors were treated with vehicle, iteration 1 peptide, iteration 5 peptide, or CTPI-2, a benchmark SLC25A1 inhibitor, with all treatments administered at equimolar doses every other day.^56^ For the adoptive-transfer model, subcutaneous B16-OVA tumors were established in C57BL/6 mice, and OT-1 CD8+ T cells (2 x 10^6) were adoptively transferred intravenously. Treatment groups included peptide plus T cells, CTPI-2 plus T cells, vehicle plus T cells, and control. Tumor volume was calculated as length x width^2 / 2 using digital calipers (Digimatic Caliper CD-15APX, Mitutoyo), and body weight and clinical signs were monitored throughout treatment.

#### METHOD DETAILS

##### Clinically informed peptide prior dataset and preprocessing

The CPP model used a shared transformer-style peptide encoder combined with abundance-aware gated fusion and cancer-type encoding. Survival, recurrence-free survival, and malignancy prediction heads were trained jointly using cross-entropy or Cox proportional-hazards-style losses according to label type. Unless otherwise specified, models were trained with an 80:10:10 train/validation/test split stratified by cancer type, AdamW optimization, a learning rate of 1e-4, batch size 32, early stopping on validation loss, and five random seeds.^75^ Favorable-outcome conditioning was used to extract label-conditioned attention matrices, which were normalized across layers and sequence positions to yield residue- and peptide-level importance scores.

Candidate peptides were filtered by length, molecular weight, net charge, pI, GRAVY, aromaticity, aliphatic index, membrane-interface compatibility, predicted half-life, solubility, safety, and permeability. Property distributions corresponding to Figure S1H were calculated using standard peptide-property functions, and candidates outside SPPS-compatible or assay-compatible ranges were excluded. Terminal caps or chemical modifications were not introduced unless explicitly specified for a labeled or control peptide. The final CPP contained 4,822 clinically anchored candidates with parent protein, cancer context, outcome label, sequence score, and physicochemical metadata.

##### GALILEO agent stack, OTAS loop and sequence editing

GALILEO implements an Observation-Thought-Action-Summary (OTAS) cognitive cycle that integrates autonomous multi-agent reasoning with iterative physical wet-lab execution. The multi-agent reasoning system comprises four in-house agent types: (1) the Architect agent, which proposes candidate peptide sequences based on the current Hypothesis Board and prior experimental outcomes; (2) the Reviewer agent, which evaluates biophysical plausibility and consistency with prior data; (3) the PI-Tech planner, which translates approved hypotheses into executable robotic protocols; and (4) adversarial Blue-team/Red-team evaluator agents, which independently argue for and against each hypothesis before physical execution. All agents were built in house and optimized for membrane-protein targeting. The Hypothesis Board serves as a shared memory substrate storing experimental outcomes, motif-performance data, and structural confidence scores. During Action, the sequence editor was constrained to amino-acid insertion, deletion and substitution on CPP-derived sequences; no unconstrained de novo generation was permitted. Each edit recorded the parent CPP peptide, operation, rationale, target hypothesis, score changes, assay command and physical outcome.

##### Peptide sequences, SPPS synthesis and HPLC-MS quality control

All peptides, including GALILEO-LRC, GALILEO-SLC, intermediate iteration peptides, fluorescent derivatives and sequence-scrambled controls, were synthesized and purified by GenScript using standard Fmoc-based solid-phase peptide synthesis, following established SPPS chemistry.^76^ Oxidative folding for cysteine-rich peptides was performed when required. Crude peptides were purified by reverse-phase HPLC system (Vanquish UHPLC, Thermo Fisher Scientific), and identity and molecular weight were confirmed by mass spectrometer (Orbitrap Exploris 120, Thermo Fisher Scientific). Only peptides with purity >95% were used for biological experiments. Peptides were dissolved in DMSO (Sigma-Aldrich) at 10 mM unless solubility required aqueous buffer, aliquoted, and stored at −80°C.

Analytical HPLC-MS quality control was performed after purification to verify peptide identity and purity. Peptides were analyzed by reverse-phase HPLC-MS system (Vanquish UHPLC coupled to Orbitrap Exploris 120, Thermo Fisher Scientific) using a C18 column (ACQUITY UPLC BEH C18, 2.1 x 50 mm, 1.7 μm; Waters) and a water/acetonitrile gradient containing 0.1% formic acid or 0.1% TFA. Peptides were accepted for biological testing when the observed mass matched the calculated mass within instrument tolerance and analytical purity was >95%. Peptides failing mass, purity, solubility or precipitation guardrails were resynthesized or excluded from the optimization loop.

##### Robotic platform and plate-based phenotypic screening

The robotic wet-lab platform integrated an automated liquid-handling system (Bravo, Agilent Technologies), a robotic arm (BenchBot, Agilent Technologies), plate-preparation modules including automated plate marker (Microplate Labeler, Agilent Technologies), tip/plate feeder (BenchCel Microplate Handler, Agilent Technologies) and labware handler (Labware MiniHub, Agilent Technologies), a microplate clamper/sealer (PlateLoc Thermal Microplate Sealer, Agilent Technologies), and an assay liquid handler for quantitative readout (MultiFlo FX Multi-Mode Dispenser, BioTek/Agilent Technologies). The assay liquid handler performed reagent dispensing and plate-based assay setup; the automated liquid-handling system (Bravo, Agilent Technologies) performed dilution, transfer and mixing steps; and the robotic arm (BenchBot, Agilent Technologies) and plate-preparation modules executed plate movement, tracking, and run supervision. Imaging and plate-reader outputs were collected using BioTek-compatible imaging or multimode plate-reader instrumentation (Cytation 5 Cell Imaging Multi-Mode Reader/Synergy H1 Hybrid Multi-Mode Reader, BioTek/Agilent Technologies). The deck layout, plate maps and machine-readable commands were generated by the GALILEO wet-lab interface module, and robotic operations were executed under GALILEO control.

Initial screening assays were performed in 96-well plates seeded with 3-5 x 10^3 cells per well. Cells were allowed to adhere overnight and treated with 10-point half-log or three-fold peptide gradients from 1 nM to 10 μM unless solubility or branch-specific confirmation assays required modified concentrations. Vehicle, PBS and sequence-scrambled peptide controls were included where appropriate. Each dose was tested in at least triplicate technical wells, and screening readouts were collected at 24, 48 and 72 h. SYTOX Green nucleic-acid stain (Thermo Fisher Scientific) was used at 1 μM for 15-30 min before imaging to quantify membrane-compromised cells. CellTiter-Glo viability reagent (Promega) was added at a 1:1 reagent-to-medium ratio, incubated for 10 min at room temperature, and read as luminescence using a multimode plate reader (Synergy H1 Hybrid Multi-Mode Reader, BioTek/Agilent Technologies). Viability and inhibition were normalized to matched vehicle or PBS controls. Branches were advanced only when tumor inhibition was observed without matched HEK293T toxicity, with a conventional advancement threshold of >=20% tumor inhibition and <=10-15% HEK293T inhibition under the same exposure window unless branch-specific rules were applied.

##### Cognitive benchmarks, motif dynamics and ABG extraction

SeqQA, FigQA, TableQA, ProtocolQA, DbQA and cloning-scenario benchmarks were evaluated using predefined question sets, rubric-based scoring and matched prompt budgets. Mean F1 was calculated as the harmonic mean of precision and recall averaged across answer categories. The complete questionnaire and rubric are provided as Supplementary Data. Motif payload dynamics were quantified across the five autonomous optimization rounds. For each round, 3-mer motif Shannon extropy was calculated as:

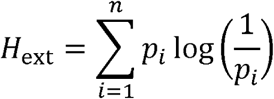

where p_i is the frequency of motif i among tested active peptides. Active motif sharing was defined by recurrent 3-mer or motif-class overlap between peptides exceeding the prespecified inhibition threshold. Enrichment of motif classes was tested by Fisher’s exact test or permutation tests with Benjamini-Hochberg correction.

##### UMAP/property analysis and Peptipedia comparison

Peptipedia peptides with explicit functional annotations were used as reference sequences.^39^ Peptides were embedded with the ESM-C protein language model, and UMAP was run with cosine distance, n_neighbors=15, min_dist=0.1, and random_state=42.^77^ GALILEO, RFdiffusion3 and BindCraft peptide outputs were scored with the PepVerse/PeptiVerse peptide-property scoring suite (GALILEO repository version) for stability, specificity, hydrophilicity, solubility, safety and permeability, using the same score versions and thresholds across all methods.

##### RFdiffusion3, BindCraft, AlphaFold3 rebuilding and benchmark scoring

RFdiffusion3 and BindCraft were used as external comparator workflows under matched computational budgets. RFdiffusion3 was run through the RosettaCommons Foundry RFdiffusion3 implementation available at analysis time, building on the RFdiffusion framework, and BindCraft was run from the public BindCraft workflow using its AlphaFold2-backpropagation/MPNN/PyRosetta design pipeline.^78^ For each target, design budgets, target structures, seed numbers, post-design filters and runtime constraints were matched where possible. Top complexes were rebuilt with AlphaFold3 using random seeds 1-8, and target-level means across the eight rebuilds were used for statistical comparisons.^27^ The target-aware design-space calculation and benchmark scoring were written as explicit mathematical definitions. For a peptide of length L, the naive sequence count was defined as:

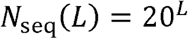

The target-conditioned sequence-property-geometry design manifold was defined as:

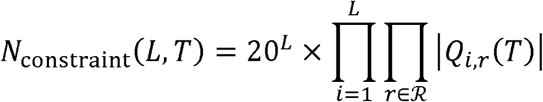

where T denotes the target, i indexes peptide positions, r indexes functional design regimes, and Q_i,r(T) denotes the target-conditioned categorical requirements for charge, hydropathy, flexibility, packing and anchoring. For known-reference targets, blocker recovery was scored by gold-reference consistency as:

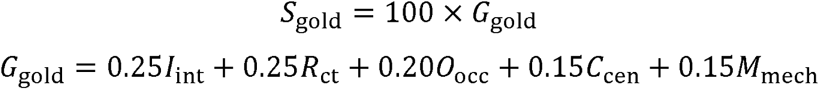

In this weighted sum, I_int, R_ct, O_occ, C_cen and M_mech denote interface recovery, contact recovery, occupancy, centroid-distance consistency and mechanism-match terms, respectively. For no-reference targets, intrinsic blocker plausibility was scored as:

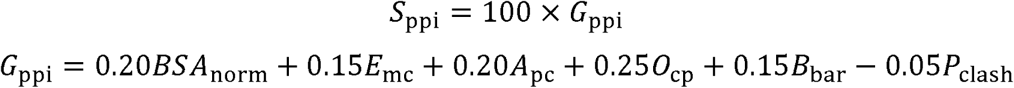

Here, BSA_norm, E_mc, A_pc, O_cp, B_bar and P_clash denote normalized buried surface area, multichain engagement, physicochemical interface compatibility, core/pore occupancy, steric-barrier formation and clash penalty terms, respectively. For plotting and statistical comparison, raw scores were clipped to the closed interval [0,100] as:

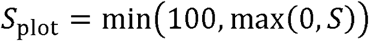

Known-reference targets were SLC7A11, CFTR, SLC25A4 and TPCN2, for which literature-supported inhibitory regions or reference ligands were available. No-reference targets were CACNA2D2, SLC16A3 and ANO1, for which selective structural reference blockers were not available under the benchmark criteria. The full 65-target benchmark included 49 SLC or SLCO transporters and 16 ion-channel or channel-associated proteins selected by CPTAC/TCGA tumor-expression or outcome association, membrane localization, transporter/channel function and lack of mature peptide-blocker coverage. This expanded set underwent virtual self-iteration only and was not synthesized or phenotyped.

##### Molecular-dynamics simulations

Membrane-embedded complexes were prepared with CHARMM-GUI and simulated with GROMACS (v2023.3).^79^ SLC25A1 complexes were embedded in a mitochondrial-mimetic lipid bilayer using CHARMM36m force-field parameters, solvated with TIP3P water, neutralized with counterions and supplemented to 150 mM NaCl.^79^ Systems were energy-minimized, equilibrated under NVT and NPT conditions with positional restraints, and simulated for a 30-ns production window at 310 K and 1 bar using a 2-fs timestep. Interface RMSF, complex RMSD and steric-barrier profiles were calculated from aligned trajectories.

##### Pan-cancer organoid membrane proteomics

Patient-derived cancer organoids from BRCA, COAD, STAD, LIHC, LUAD and PAAD were used for the pan-cancer membrane-proteomic atlas. Membrane protein enrichment was performed using the Mem-PER Plus Membrane Protein Extraction Kit (Thermo Fisher Scientific). Proteins were reduced with 10 mM DTT at 56°C for 30 min, alkylated with 55 mM iodoacetamide for 30 min in the dark, and digested with trypsin at a 1:50 enzyme-to-protein ratio at 37°C for 16 h. Peptides were desalted using C18 StageTips (Empore C18 extraction disks, 3M) and analyzed by nanoLC-MS/MS system composed of a Vanquish UHPLC system (Thermo Fisher Scientific) coupled to an Orbitrap Exploris 120 mass spectrometer (Thermo Fisher Scientific) under a standard data-dependent acquisition workflow. Peptides were separated on a nanoLC reversed-phase C18 column (EASY-Spray PepMap RSLC C18, 75 μm x 50 cm, 2 μm; Thermo Fisher Scientific) with a water/acetonitrile gradient containing 0.1% formic acid. Raw data were processed using MaxQuant (v2.1.4) against the UniProt human database.

##### LRRC8C campaign methods

LRRC8C was nominated from a multi-cancer organoid membrane-proteomic atlas containing BRCA, COAD, STAD, LIHC, LUNG and PAAD organoids, comprising 104 organoid samples. The agent identified 35 recurrent ion-channel candidates and ranked them by detection rate, tumor-versus-normal expression context, druggable-pore plausibility, immune-context relevance, literature evidence and unmet inhibitor need. LRRC8C expression and exhaustion scores were evaluated across 22 CancerSCEM cancer types using standard cell-type annotations and an exhaustion score defined by detected expression of PDCD1, HAVCR2, LAG3, TIGIT, CTLA4, TOX and ENTPD1 in CD8+ T cells.^80^

Whole-cell patch-clamp recordings were performed in HEK293T cells transiently expressing defined LRRC8 subunit combinations, following the general whole-cell recording strategy and LRRC8/VRAC functional principles.^41^ HEK293T cells were co-transfected with LRRC8A plus LRRC8C, LRRC8B, LRRC8D or LRRC8E plasmids at a 1:1 ratio. Recordings were conducted 24-48 h after transfection using a patch-clamp amplifier (Axopatch 200B, Molecular Devices) and electrophysiology acquisition software (pClamp, Molecular Devices). Patch pipettes with 3-5 MΩ resistance were used, and cells were held at −70 mV. Hypotonic solution (220 mOsm kg-1) was applied to activate VRAC currents. GALILEO-LRC was applied acutely after current activation at 3.9-31.2 μM. Currents were sampled at 2 kHz, filtered at 1 kHz and analyzed using electrophysiology analysis software (Clampfit, Molecular Devices) and GraphPad Prism (v10, GraphPad Software).

LRRC8C-high and LRRC8C-low breast cancer organoids were treated for 72 h with PBS, sequence-scrambled peptide or GALILEO-LRC. Organoid morphology and area distributions were quantified from five randomly selected fields per condition using ImageJ.^81^ For multicancer panel assays, Huh-7, HepG2, Hep3B, SNU-182, HuCCT1, A549, PC9, MKN45, HeLa, HCT116 and K562 cells were treated for 72 h and analyzed by CellTiter-Glo viability assay (Promega) or imaging-based growth inhibition using an imaging system or multimode plate reader (Synergy H1 Hybrid Multi-Mode Reader, BioTek/Agilent Technologies). Jurkat or primary T-cell viability controls were treated under matched peptide conditions to evaluate direct T-cell toxicity.

For LRRC8C metabolomics, Huh-7 cells were treated with GALILEO-LRC, PBS vehicle or sequence-scrambled peptide controls. Metabolites were extracted from cell pellets using pre-chilled methanol/acetonitrile-based solvent systems with repeated freeze-thaw and sonication steps, followed by protein precipitation at −40°C and centrifugation. LC-MS/MS analysis was performed using a UHPLC system (Vanquish UHPLC, Thermo Fisher Scientific) coupled to a mass spectrometer (Orbitrap Exploris 120, Thermo Fisher Scientific). Polar metabolites were separated on a hydrophilic-interaction LC column (ACQUITY UPLC BEH Amide, 2.1 x 50 mm, 1.7 μm; Waters). The mobile phase consisted of 25 mM ammonium acetate and ammonium hydroxide in water (A) and acetonitrile (B), and data were acquired in both positive and negative ion modes. Raw data were converted to mzXML using ProteoWizard/MSConvert (v3.0, ProteoWizard) and processed with XCMS-based pipelines for peak detection, alignment and integration.^82^ Metabolites were annotated using in-house databases with MSI classification criteria, normalized using total ion current (TIC), and analyzed by t test (P < 0.05) and VIP > 1 from OPLS-DA. Pathway enrichment was performed using KEGG and MetaboAnalyst.^83^ For RNA-seq, total RNA was extracted using the RNeasy kit (Qiagen), assessed with TapeStation system (4200 TapeStation, Agilent Technologies) and fluorometer/spectrophotometer systems (Qubit 4 Fluorometer and NanoDrop One, Thermo Fisher Scientific), and prepared using the Illumina Stranded mRNA Prep, Ligation kit (Illumina). Libraries were sequenced on a sequencing platform (DNBSEQ-T7, MGI Tech) with paired-end 150-bp reads. Raw reads were processed using trim_galore (v0.6.4; wrapper around Cutadapt), rRNA removal was performed with bowtie2 (v2.4.2), clean reads were aligned to GRCh38.104 using HISAT2 (v2.2.0), and gene-level counts were generated with htseq-count (v0.13.5).^84^ Differential expression analysis was performed with edgeR, and GO/KEGG enrichment was performed with clusterProfiler using Fisher’s exact test.^85^ FPKM values and log2 transformation were used for downstream visualization and PCA analysis.

ROS was measured using a ROS-sensitive fluorescent probe (DCFH-DA; Reactive Oxygen Species Assay Kit, Beyotime, Cat# S0033S) at 5-10 μM for 30 min before flow-cytometry analysis on a flow cytometer (CytoFLEX S, Beckman Coulter). TMRE mitochondrial membrane potential dye (Thermo Fisher Scientific, Cat# T669) was used at 50-100 nM for 20-30 min to quantify mitochondrial membrane potential. cGAMP was measured in paired extracellular and intracellular fractions using a commercial ELISA kit (cGAMP ELISA kit, Cayman Chemical, Cat# 501700); intracellular values were normalized to 10^6 cells or total protein. STING activation was assessed by immunoblotting in HepG2 cells after 3.9, 7.81, 15.62 and 31.2 μM GALILEO-LRC treatment using phospho-STING (Ser366) antibody (Cell Signaling Technology, Cat# 19781), total STING antibody (Cell Signaling Technology, Cat# 13647), and alpha-tubulin antibody (Cell Signaling Technology, Cat# 2144). HepG2 was used as an orthogonal hepatocellular carcinoma context with robust STING immunoblot signal.

LRRC8C target dependence was tested by siRNA-mediated LRRC8C silencing in Huh-7 cells. Cells were transfected with LRRC8C-targeting or negative-control siRNAs using Lipofectamine RNAiMAX (Thermo Fisher Scientific), assayed 48-72 h after transfection, and verified by qPCR using a real-time PCR system (QuantStudio 5, Applied Biosystems/Thermo Fisher Scientific). JC-1 aggregate/monomer ratios were quantified by flow cytometer (CytoFLEX S, Beckman Coulter) after GALILEO-LRC exposure.

For Huh-7/T-cell cocultures, Huh-7 cells were seeded one day before adding HLA-compatible primary human T cells at a conventional E:T ratio of 5:1. Cultures were treated with control, diABZI or GALILEO-LRC for 48h. CD107a antibody was added during the assay, with monensin or brefeldin A used during the final 4-6 h when required. Cells were stained for CD3, CD8, CD107a and viability dyes and analyzed by flow cytometer (CytoFLEX S, Beckman Coulter). For LIHC-PDTF immune assays, treated fragments were stained for GZMB by immunohistochemistry or dissociated for CD3/CD69 flow cytometry. Isolated T cells cultured without tumor fragments were included to test direct peptide-driven activation or toxicity.

For LRRC8C in vivo testing, female C57BL/6 mice (7 weeks old) were subcutaneously inoculated with 2 x 10^6 Hepa1-6 cells. When tumors reached 50-100 mm3, mice were randomized to vehicle, GALILEO-LRC peptide (40 μg per dose) or diABZI (40 μg per dose).^86^ Treatments were administered every 3 days for four doses, and tumor volume was measured every 2 days. Endpoint samples were analyzed by flow cytometer (CytoFLEX S, Beckman Coulter) for Annexin V/FVD780 T-cell death and by tumor-volume comparison.

##### SLC25A1 campaign methods

The SLC25A1 target-nomination workflow integrated TCGA, GTEx, GEO, CPTAC, immune-infiltration analysis, literature screening and translational-gap assessment.^34,87^ Candidate SLC targets were ranked across functional uniqueness, pan-cancer overexpression, prognostic relevance, druggability potential, stemness and drug-resistance association, and immune modulation. CIBERSORT deconvolution was run with standard LM22 immune-cell signatures, and partial Spearman correlations were calculated while controlling for tumor purity or cancer-type covariates where appropriate. SLC25A1 was prioritized over SLC7A11, SLC16A1, SLC25A51, SLC1A5 and SLC2A1.

CTPI-2 was used as a matched-molar benchmark SLC25A1 small-molecule comparator. CTPI-2 stocks were prepared in DMSO and diluted into culture medium immediately before treatment with final DMSO <=0.1%. Unless stated otherwise, direct GALILEO-SLC and CTPI-2 comparisons used identical molar exposure. GALILEO-SLC mitochondrial localization was assessed using 5-FAM-Ahx-labeled peptide and MitoTracker Red CMXRos staining (Thermo Fisher Scientific, Cat# M7512), imaged by confocal microscope (LSM 880, Zeiss) after 30-60 min of staining. Colocalization was assessed qualitatively and, where quantified, by Pearson correlation coefficient in ImageJ/Fiji.

SLC25A1 silencing was performed using siRNA-mediated depletion in Huh-7 cells, followed by JC-1 analysis after GALILEO-SLC treatment. Cells were transfected with SLC25A1-targeting or negative-control siRNA using Lipofectamine RNAiMAX (Thermo Fisher Scientific), treated 48-72 h later and verified by qPCR using a real-time PCR system (QuantStudio 5, Applied Biosystems/Thermo Fisher Scientific) or immunoblotting. BCO9 organoids were treated with GALILEO-SLC or equimolar CTPI-2 for 72 h, and organoid area distributions were quantified from five randomly selected brightfield fields per condition. Dose-response curves were fit by nonlinear regression to estimate IC50.

For BODIPY/TMRE/acetate rescue, Huh-7 cells were treated for 72 h with GALILEO-SLC, equimolar CTPI-2, vehicle or acetate rescue. Sodium acetate was added at a conventional rescue concentration of 2-5 mM at the time of peptide treatment unless otherwise specified. Lipid droplets were stained with BODIPY 493/503 (Thermo Fisher Scientific, Cat# D3922), mitochondria were stained with TMRE mitochondrial membrane potential dye (Thermo Fisher Scientific, Cat# T669), nuclei were stained with Hoechst 33342 (Thermo Fisher Scientific, Cat# H3570), and images were quantified with ImageJ/Fiji macros for lipid-droplet number per cell and mean TMRE fluorescence intensity.

SLC25A1 metabolomics compared iteration 1 and iteration 5 peptides at matched inhibitory effect. Huh-7 cells were extracted using the same cold methanol/acetonitrile-based workflow described above. LC-MS/MS was performed on a UHPLC system (Vanquish UHPLC, Thermo Fisher Scientific) coupled to a mass spectrometer (Orbitrap Exploris 120, Thermo Fisher Scientific), and polar metabolites were separated on a hydrophilic-interaction LC column (ACQUITY UPLC BEH Amide, 2.1 x 50 mm, 1.7 μm; Waters). Raw data were converted to mzXML with ProteoWizard, processed with XCMS, normalized by TIC and analyzed by t test (P < 0.05) with VIP > 1 from OPLS-DA.^88^ Pathway enrichment used KEGG and MetaboAnalyst.^83^ Transcriptomic profiling compared iteration 1 and iteration 5 peptides against matched untreated controls. RNA extraction, quality control, library preparation, DNBSEQ-T7 sequencing, read processing, alignment, htseq-count quantification, edgeR differential expression, and clusterProfiler enrichment followed the transcriptomics workflow described above. Autophagic flux was quantified by LC3/p62 immunostaining, LC3B immunoblot or a validated autophagic-flux reporter as available. AMPK activation was assessed by immunoblotting with phospho-AMPKα (Thr172) antibody (Cell Signaling Technology, Cat# 2535) and AMPKα antibody (Cell Signaling Technology, Cat# 2532) after 48 h treatment with 58.6, 234.4, 937.5 and 3,750 nM GALILEO-SLC.

Extracellular pH was measured in Huh-7 cells, BCO breast cancer organoids and LIHC-PDTF cultures using a calibrated micro-pH electrode (InLab Micro Pro-ISM, Mettler Toledo) or plate-compatible pH system (SensorDish Reader SDR with HydroDish pH sensor plates, PreSens) at the same post-treatment time point used in Figure 6D. Culture medium and CO2 exposure were standardized across groups before measurement. Seahorse ECAR was measured in Huh-7 cells using a Seahorse XF analyzer (Seahorse XFe96 Analyzer, Agilent Technologies) with an XF glycolytic stress test. Cells were seeded in Seahorse XF plates (Agilent Technologies) one day before assay, equilibrated in bicarbonate-free Seahorse assay medium (Agilent Technologies) supplemented with 2 mM glutamine, and sequentially injected with glucose (10 mM), oligomycin (1 μM) and 2-deoxyglucose (50 mM). ECAR was normalized to cell number or protein content.

For SLC25A1 in vivo testing, C57BL/6 mice bearing Hepa1-6 tumors were treated with vehicle, iteration 1 peptide, iteration 5 peptide or CTPI-2. All treatments were administered at equimolar doses every other day. In the adoptive-transfer model, subcutaneous B16-OVA tumors were established in C57BL/6 mice, and OT-1 CD8+ T cells (1 x 10^6) were adoptively transferred intravenously. Treatment groups included peptide plus T cells, CTPI-2 plus T cells, vehicle plus T cells and control. Tumor growth was monitored over time, and endpoint tumors were analyzed by CD69/CD8 flow cytometry using a flow cytometer (CytoFLEX S, Beckman Coulter) or GZMB immunohistochemistry.

##### Immunohistochemistry and flow cytometry

For GZMB immunohistochemistry, tissues were fixed in 4% paraformaldehyde or formalin, embedded in paraffin or OCT according to tissue type, sectioned at 4-5 μm using a microtome (RM2235, Leica Biosystems) or cryostat (CM1950, Leica Biosystems), subjected to antigen retrieval in citrate or EDTA buffer, blocked, incubated with anti-GZMB antibody overnight at 4°C, and developed with HRP-DAB or fluorescent secondary detection. Five random fields per sample were quantified by ImageJ (NIH; Schneider et al., 2012) or QuPath (v0.4.3, QuPath developers) using a fixed threshold, and the number of GZMB-positive cells was normalized to 10^6 pixels or tissue area.^89^

All flow-cytometry data were acquired on a flow cytometer (CytoFLEX S, Beckman Coulter) and analyzed using FlowJo software (v10, BD Biosciences). Flow-cytometry panels included CD3, CD8, CD69, CD107a, Annexin V, FVD780 viability dye, JC-1, DCFH-DA ROS probe, TMRE and other mitochondrial-potential dyes as required by each assay. ROS, mitochondrial membrane potential and immunogenic-cell-death assays were performed using standard fluorescent probes (DCFH-DA, TMRE, JC-1) and Annexin V/FVD staining according to manufacturer protocols. Compensation was performed using single-stain controls, dead cells and debris were excluded, and at least 10,000 live events were collected per sample when feasible.

#### QUANTIFICATION AND STATISTICAL ANALYSIS

All statistical analyses were performed using GraphPad Prism (v10, GraphPad Software), R (v4.3.1, R Foundation for Statistical Computing; R Core Team, 2023) and the indicated Python packages (Python v3.11.2, Python Software Foundation). Data are presented as mean ± s.d. or mean ± s.e.m. as indicated in figure legends, with animal tumor-growth panels shown as mean ± s.e.m. unless otherwise specified. Statistical tests included unpaired or paired two-tailed Student’s t tests, Wilcoxon tests for non-normally distributed or rank-based comparisons, one-way or two-way ANOVA with post-hoc corrections for multi-group comparisons, and repeated-measures two-way ANOVA or mixed-effects models for tumor-growth curves. IC50 values were estimated by four-parameter nonlinear regression. Significance was defined as P < 0.05 unless otherwise specified and reported as ns, not significant; *P < 0.05; **P < 0.01; ***P < 0.001; ****P < 0.0001.

## SUPPLEMENTAL TABLE LEGENDS

**TableS1. Comparative capability matrix for GALILEO and representative AI-scientist systems, related to Figure 1D**.

**Table S2. GALILEO closed-loop optimization event log and hypothesis-board audit, related to Figures 1, 2, and 5**.

**Table S3. Literature-supported ion-channel target evidence for GALILEO target nomination, related to Figure 3A**.

## SUPPLEMENTAL VIDEO LEGENDS

**Video S1. The video features GALILEO, an embodied AI scientist that combines AI-agent, robotics and autonomous wet-lab validation to accelerate the discovery of therapeutic interventions, related to Figure 1B**.

## Notes

### Competing Interest Statement

The authors have declared no competing interest.

