## Supplemental figure for "GALILEO: Embodied AI scientist for autonomous therapeutic discovery in dynamic membrane systems"

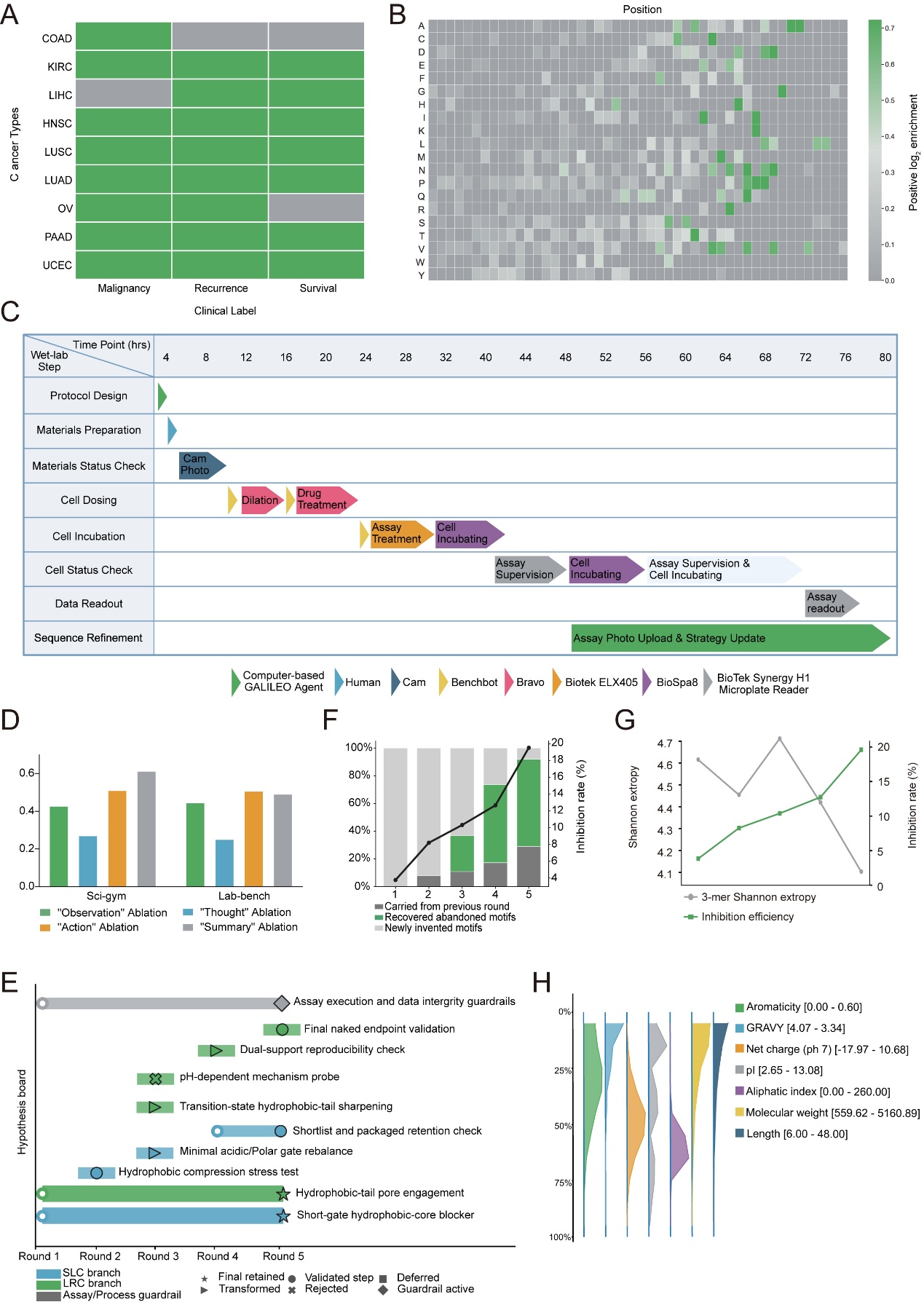

**Figure S1. CPP construction, autonomous wet-lab scheduling, and motif-evolution audit.**

(A) CPTAC clinical-label coverage across cancer types and outcome labels used for CPP training.

(B) Unified amino acid attention matrix generated by normalizing cancer-specific attention maps across model layers and sequence positions.

(C) Gantt-style schedule of one autonomous wet-laboratory cycle, including protocol design, material preparation, cell dosing, incubation, imaging, assay treatment, data readout, and sequence refinement over the indicated time window.

(D) Component ablation analysis on SciGym and LAB-Bench comparing Observation, Thought, Action, and Summary module removal.

(E) Hypothesis-board evolution for SLC and LRC branches across five rounds, including assay/process guardrails, validated steps, rejected hypotheses, deferred hypotheses, and final retained mechanisms.

(F) Iteration-wise contributions of motifs carried from the previous round, recovered abandoned motifs, and newly invented motifs, overlaid with inhibition rate.

(G) 3-mer Shannon extropy and inhibition efficiency across five autonomous optimization rounds.

(H) Physicochemical-property distributions of 4,822 CPP peptides, including length, molecular weight, net charge at pH 7, pI, GRAVY, aromaticity, and aliphatic index.

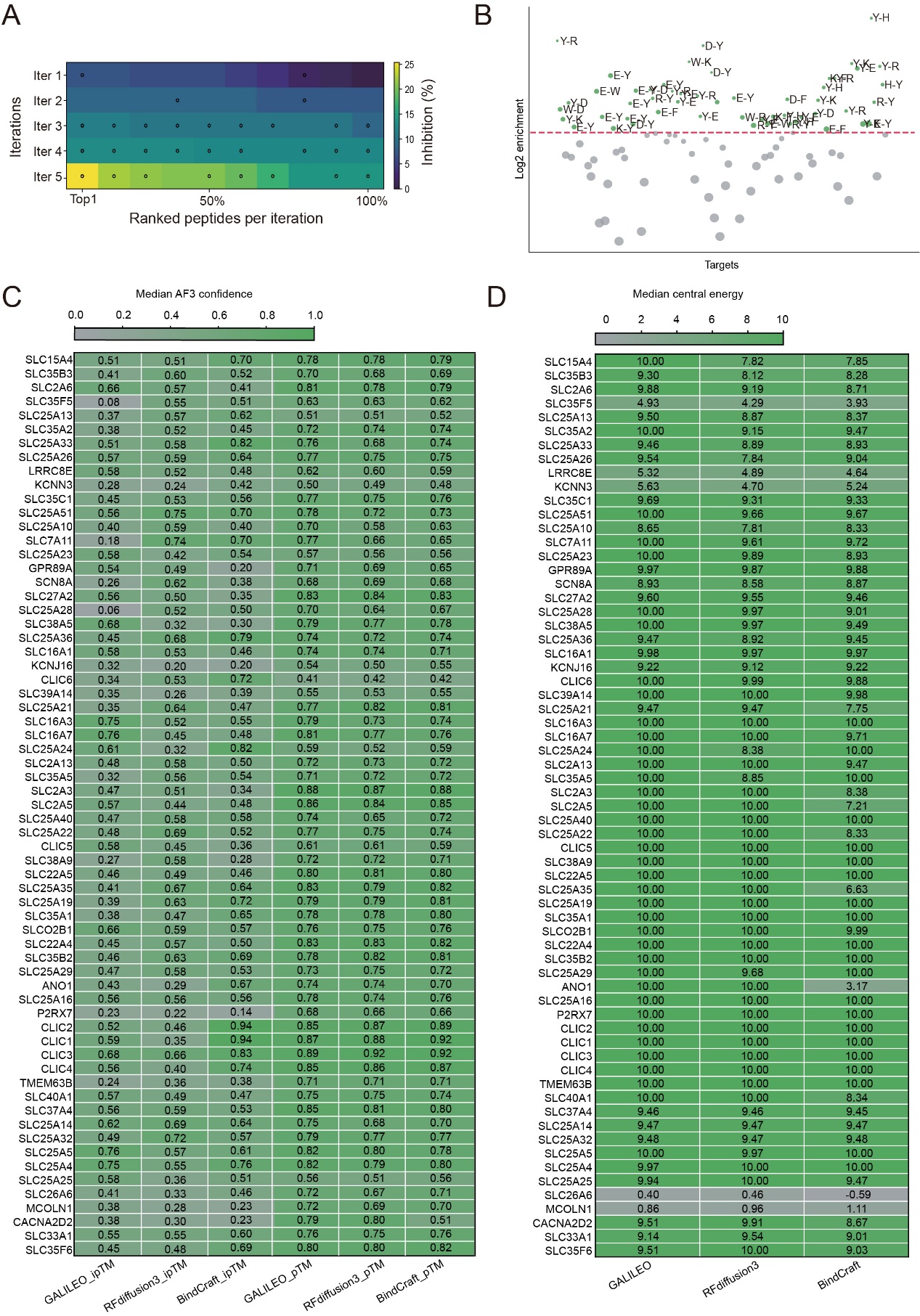

**Figure S2. Benchmark-scale virtual iteration and Amphiphilic Balance Grammar features.**

(A) Heatmap of ranked peptide inhibition across five autonomous iterations. Dots indicate peptides that inhibited at least two tumor cell lines among Huh-7, SNU-182, HCCLM3, and Hep3B while sparing HEK293T cells.

(B) Log2 enrichment of amino acid pair features defining the ABG grammar across target contexts. Positive enrichment indicates motif-pairing features retained by GALILEO after physical or virtual iteration.

(C) Median AlphaFold3 confidence metrics (ipTM and pTM) for GALILEO, RFdiffusion3, and BindCraft peptide-target complexes across 65 benchmark proteins after uniform AlphaFold3 rebuilding. The benchmark includes 49 SLC or SLCO transporters and 16 ion-channel or channel-associated proteins.

(D) Median central steric-barrier energy for GALILEO, RFdiffusion3, and BindCraft peptide-target complexes across benchmark proteins.

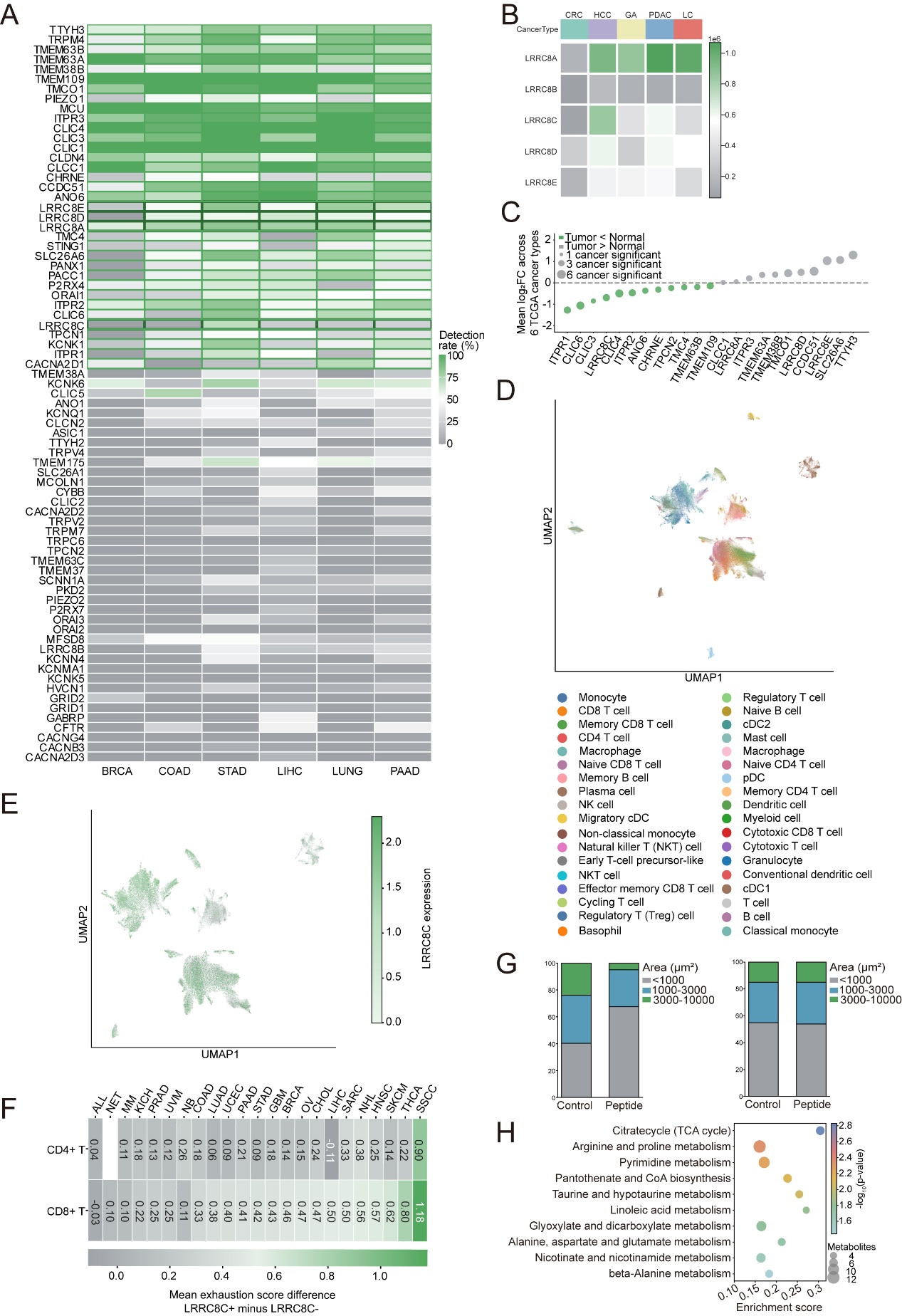

**Figure S3. LRRC8C target nomination and orthogonal validation.**

(A) Detection-rate heatmap of recurrent ion channels across BRCA, COAD, STAD, LIHC, LUNG, and PAAD in the multi-cancer organoid membrane-proteomic atlas, ranked by cross-cancer prevalence.

(B) Heatmap of LRRC8 family abundance across CRC, HCC, gastric adenocarcinoma (GA), pancreatic ductal adenocarcinoma (PDAC), and lung cancer (LC) organoid datasets.

(C) Mean log2 fold-change across six TCGA cancer types for prioritized ion-channel candidates; point size indicates the number of cancer types with significant differential expression.

(D) Batch-integrated immune-cell UMAP generated from ALL, NET, MM, KICH, PRAD, UVM, NB, COAD, LUAD, UCEC, PAAD, STAD, GBM, BRCA, OV, CHOL, LIHC, SARC, NHL, HNSC, SKCM, THCA, and SSCC single-cell datasets.

(E) LRRC8C expression projected onto immune-cell clusters from D.

(F) Cancer-type-specific exhaustion-score difference between LRRC8C-positive and LRRC8C-negative CD8+ T cells.

(G) Area-distribution analysis of LRRC8C-high BCO9 organoids (left) and LRRC8C-low BCO13 organoids (right) after control or GALILEO-LRC treatment.

(H) Pathway enrichment of untargeted metabolomic changes after LRRC8C blockade, highlighting taurine/hypotaurine metabolism together with TCA-cycle and amino-acid-associated programs.

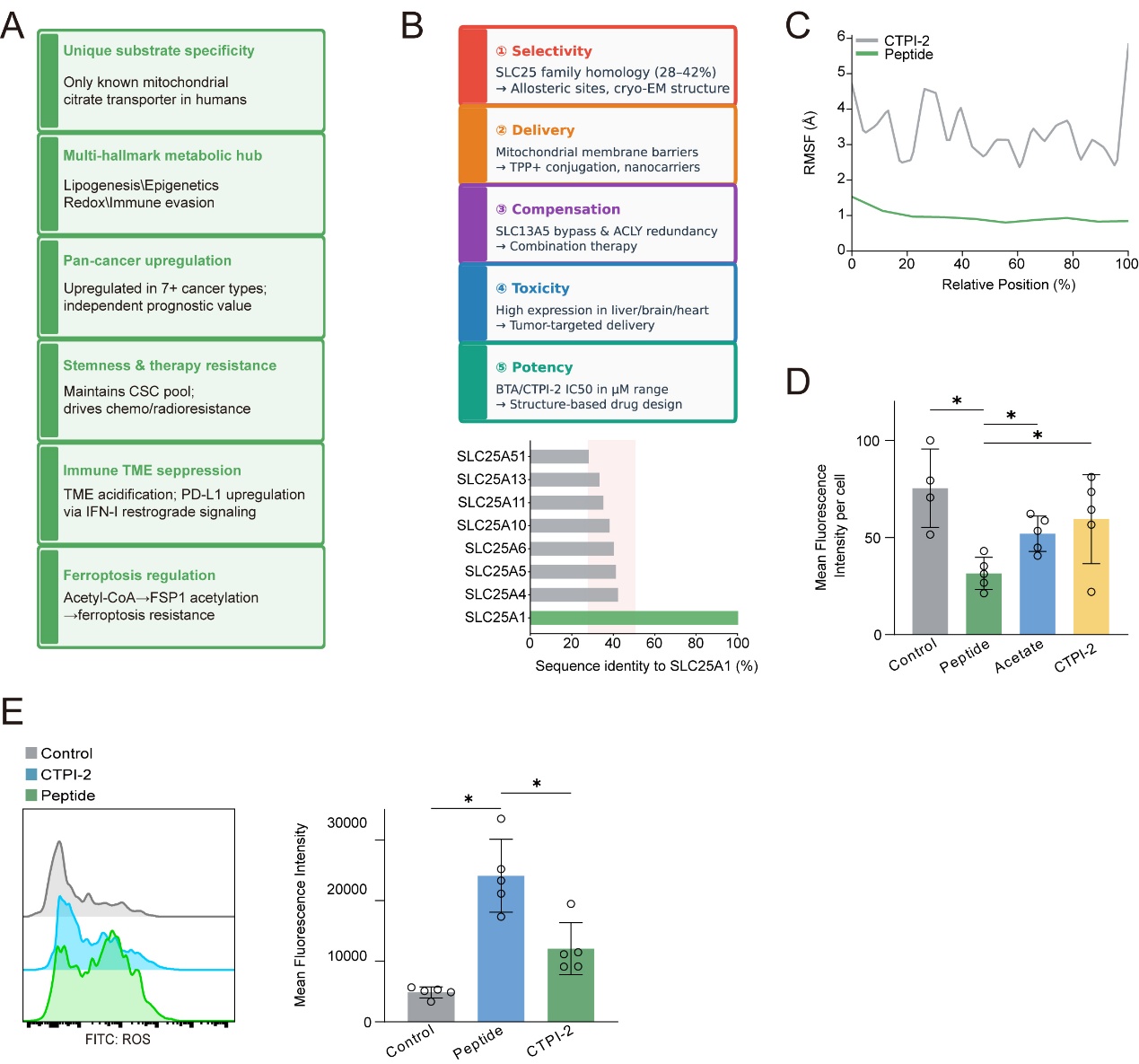

**Figure S4. Translational rationale for SLC25A1 targeting and orthogonal validation of GALILEO-SLC-induced mitochondrial stress.**

(A) Biological and translational features supporting SLC25A1 as a pan-cancer target, including unique substrate specificity, metabolic-hub function, broad overexpression, stemness/drug-resistance relevance, immune-TME suppression, and ferroptosis regulation.

(B) Major limitations of current small-molecule strategies for SLC25A1 inhibition, together with sequence-identity relationships across the SLC25 family.

(C) Molecular-dynamics RMSF comparing GALILEO-SLC with CTPI-2 across aligned sequence position in the same CHARMM-GUI-generated mitochondrial membrane system used in Figure 5G.

(D) Orthogonal TMRE red-channel quantification per cell after control, GALILEO-SLC, acetate, or CTPI-2 treatment, complementing Figure 5L.

(E) Orthogonal ROS measurement comparing control, GALILEO-SLC, and CTPI-2; left, representative FITC histograms; right, quantification of mean fluorescence intensity. Data are mean ± s.d. Individual points denote independent experiments. P values were determined by unpaired two-tailed Student's t test. See also Figure 5.
